# Predicting Permeation of Compounds across the Outer Membrane of *P. aeruginosa* Using Molecular Descriptors: Advantages and Limitations

**DOI:** 10.1101/2023.09.02.555818

**Authors:** P. D. Manrique, I. V. Leus, C. A. Ĺopez, J. Mehla, G. Malloci, S. Gervasoni, A. V. Vargiu, R. Kinthada, L. Herndon, N. W. Hengartner, J. K. Walker, V. V. Rybenkov, P. Ruggerone, H. I. Zgurskaya, S. Gnanakaran

**Author notes:** Correspondence (PDM), (SG).

## Abstract

**The ability of Gram-negative pathogens to adapt and protect themselves against antibiotics is a growing threat to public health. The low permeability of the outer membrane (OM) in combination with effective multidrug efflux pumps, constitute the two main antibiotic resistance mechanisms. Though much efforts have been devoted to discover new antibiotics that can bypass these defense mechanisms, no new antibiotic classes have been introduced into clinics in the last 35 years. Models that identify specific descriptors of molecular properties and predict the likelihood that a given compound is capable of successfully permeate the OM and inhibit bacterial growth while avoiding efflux could facilitate the discovery of novel classes of antibiotics. Here we evaluate 174 molecular descriptors of 1260 antimicrobial compounds and study their correlations with antibacterial activity in Gram-negative *Pseudomonas aeruginosa*. While part of these descriptors are computed using traditional approaches based on the physicochemical properties intrinsic to the compounds, ensemble docking and all-atom molecular dynamics (MD) simulations are used to derive additional bacterium-specific mechanistic properties. Descriptors of compound permeation across the OM were calculated using all-atom MD simulations of the compounds in different subregions of the OM model. Descriptors of interactions with efflux pumps were calculated from ensemble docking of compounds targeting specific binding pockets of MexB, the major efflux transporter of *P. aeruginosa*. Using these descriptors and the measured antibacterial inhibitory concentrations of compounds, we design and implement a statistical protocol to identify a subset of the molecular properties that are predictive of whether a given compound is a strong or weak permeator across the Gram-negative OM. Our results indicate that 88.4% of the compounds that show measurable antibacterial activity, follow very consistent rules of permeation, which highlight the critical role that the interaction between the compound and the OM have at predicting permeation. The remaining 11.6% of the compounds, although less predictive, are characterized by distinctive structural markers that can be used to minimize classification errors. An implementation of the permeation rules and the structural markers uncovered in our study is shown, and it demonstrates the accuracy of our approach in a set of previously unseen compounds. Taken together, our analysis sheds new light on the key molecular properties that drug candidates should have in order to be effective at OM permeation/inhibition of *P. aeruginosa*, and opens the gate to similar data-driven studies in other Gram-negative pathogens.**

## 1 Introduction

The emerging antibiotic resistance crises are driven by the indiscriminate use of existing antibiotics and the lagging discovery of new antibiotics [1, 2]. This has fueled rise of bacterial resistance at unprecedented rates. According to the World Health Organization priority list, all three pathogens classified as critical (its most urgent category) are Gram-negative [3–13]. Yet no new major class of antibiotics has been approved to treat infections caused by this group of organisms since 1962 [10, 14]. Therefore, there is a critical need to find effective ways to bypass the biological and chemical challenges that hamper the discovery of new and effective antibacterial treatments.

The major determinants of resistance in Gram-negative bacteria are (1) the low permeability barrier of the outer membrane (OM) that hinders diffusion of drug molecules across the membrane, and (2) the action of multidrug efflux pumps that expel drugs and other noxious compounds from the cytoplasm and periplasm back into the extracellular environment [4–8, 15]. The synergistic relationship between slow permeation and efflux effectively prevents intracellular accumulation of antibiotic to reach critical concentration levels that inhibit bacteria growth. Mathematical modeling efforts have been able to quantify critical aspects of single-cell in/out flux dynamics [16–18] and their implications at the colony level [19,20]. However, the large complexity and diversity of the interactions occurring between the drugs and the determinants of antibiotic resistance at molecular scale makes it harder to develop predictive models for drug permeation, efflux avoidance and antibacterial activity. Therefore, there is a need to incorporate detailed molecular determinants from computational approaches that probe the bacterium-specific molecular-level interaction profiles.

Molecular dynamics (MD) simulation has emerged as a useful tool to provide mechanistic understanding of the structure-function relationship of complex macromolecules and how they behave and interact with particular environments at the molecular scale. In addition, MD is able to offer spatio-temporal information that can fill the gap between experimental resolution and modelling limitations. MD has been successfully applied in the field of drug discovery [21–24], and in particular at providing detailed information of the molecular structures responsible for the low permeability of Gram Negative OM [25–29] and on the drug trafficking through efflux pumps [30]. Furthermore, it allows us to explore the complexity of the chemical environment by quantifying the interactions between a wide spectrum of compounds with specific proteins and compartments of a bacterial cell such as the OM and an efflux pump. The output of these MD simulations are often long multivariate time series describing the position of every atom over time, which are generally challenging to analyze. Traditional statistical techniques [31–34], network theory [35, 36], and artificial intelligence [37] are among the most implemented and promising quantitative tools to help unravel complex patterns in these large multidimensional datasets.

In *P. aeruginosa*, the low permeability of the OM is mostly attributed to its particular composition. A combination of highly anionic Lipopolysaccharide (LPS) molecules, tightly complexed with divalent cations makes this membrane an almost impenetrable shield [7, 25, 38]. A single LPS molecule provides distinctive chemical environments across the OM of the Gram-negative bacteria: 1) Long carbohydrate-enriched regions massively shield the exposure of the membrane towards the extracellular space; 2) phosphates and ionizable chemical groups are highly repellent to hydrophobic molecules and 3) a sheet of divalent cations provides strong coordination among the LPS of the outer layer in the OM. Thus, extracting the molecular determinants of the process governing the passive diffusion of molecules across this layer would be of tremendous aid in the design of new antibiotics. In order to achieve this, we have carried out massive MD calculations at atomic level, extracting specific properties during the assisted translocation of hundreds of compounds across the OM of *P. aeruginosa*. For each compound we have computed 35 permeability descriptors, which are extracted by instantiating MD trajectories from seven distinctive regions of the OM [37]. These regions were selected in order to have a full representation of all the chemical environments which directly affects the diffusion process. The approach is iteratively repeated until the entire set of compounds is fully covered. The permeability descriptors encompass a set of physical parameters which can directly impact the efficiency of molecular translocation: molecular interaction energy with the surrounding environment (Δh), number of hydrogen bonds with surrounding (HB), molecular lateral mean squared displacement (Δxy) and molecular entropy (Δs).

The major efflux pump of *P. aeruginosa* that contributes to clinical antibiotic resistance is MexAB-OprM, which extends across the inner and outer membrane aiding the organism to expel toxins from the intracellular and periplasmic region, directly into the extracellular space. [39–41]. In this complex, MexB is a homotrimeric protein embedded into the inner membrane and belonging to the Resistance Nodulation cell Division (RND) superfamily. It is in charge of recognition, binding, and transport of diverse substrates [41–43], and it constitutes the main barrier that any compound in the intracellular region needs to overcome. Each monomer of the MexB trimer adopts three different conformations enabling access (A), binding (B), and extrusion (C) of substrates [44,45]. We quantify the interactions between each of the studied compounds and MexB via ensemble docking calculations, from which we collected all docking poses (600 per compound), average affinity binding, and identified the contacts made by each compound to every MexB residue. From the list of contacts, we selected a subset of residues of the access of the Loose monomer (AP) and deep (DP) of the Tight monomer substrate binding pockets based on known crystallographic data for AcrB from *E. coli*, homologous to MexB from *P. aeruginosa* [44, 46]. These residues are generally considered to line/define the two pockets and are relevant for recognition/binding of compounds [45, 47]. Some of them, in particular the PHE residues of the hydrophobic trap inside the DP, were found to be key for the interaction of the transporter with inhibitors [48, 49]. Our computational analysis of MexB yielded 66 docking descriptors for each compound [37].

In this paper, we analyze the growth inhibitory activities of a unique library of 1260 antimicrobial molecules belonging to several structural classes of compounds including known antibiotics and efflux pump inhibitors [Fig. 1(a)]. The antibacterial activities are measured in strategically designed strains of *P. aeruginosa* [Fig. 1(b)] that can isolate the effects of permeation and efflux avoidance [50]. We then use the antibacterial activity data to identify correlations with a large set of computationally-derived mechanistic descriptors described above [Fig. 1(c)]. These properties are subsequently characterized by means of their ranked correlations along with a hierarchical clustering algorithm to establish similarity relationships (linear and non-linear) among them. The resultant clusters are used as input parameters of a statistical model that, using the experimental 50% inhibition concentration (IC_50_) data, identifies non-trivial relationships between different sets of descriptors and their ability to predict bacterial permeation. Our analysis identifies an optimal subset of nine relevant clusters containing the mechanistic markers yielding prediction accuracy scores of up to 96%. Our results highlight the role of the permeation descriptors quantifying the interactions between the compounds and the OM surface, the LPS lipid-A and oligosaccharide core 2 sub-regions of the OM. These features, combined with intrinsic properties of the compound like the hydrophobic surface area and the Randic index, show high correlations with permeation and growth inhibition information for a specified range of descriptor values. Our findings shed a new light into which specific molecular interactions are responsible for OM penetration and hindering of bacterial growth. Our approach and conclusions can impact the design of a new generation of antimicrobials.

**Figure 1:**
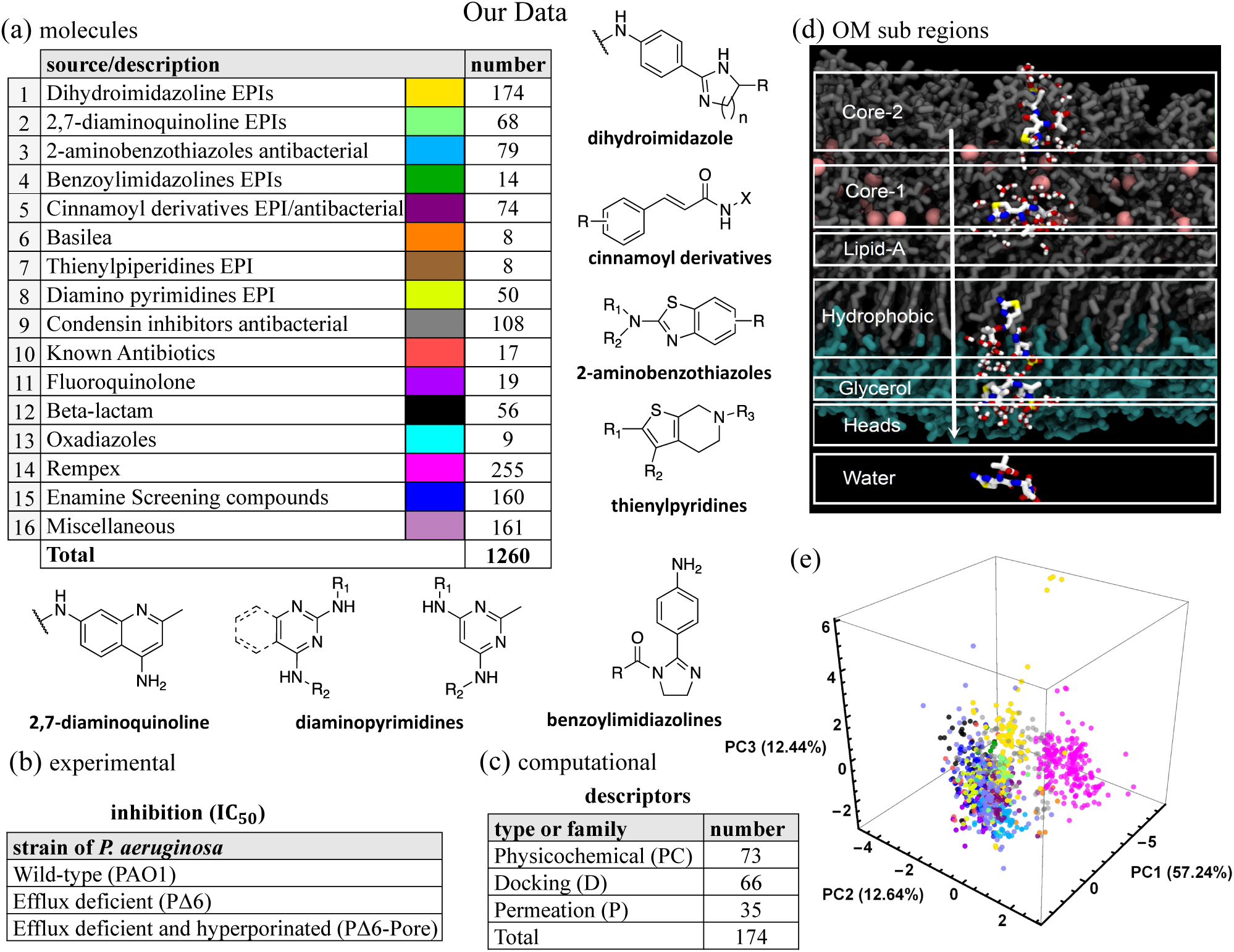
Our data is comprised by (a) 1260 antimicrobial molecules classified into 16 distinct structural chemotypes as listed (top left), and some examples are shown in the central panels. (b) Each compound is characterized by its antimicrobial activity in three strains of *P. aeruginosa* by means of the 50% inhibitory concentration (IC_50_). (c) In addition, the molecules in (a) are further characterized by 174 computationally-derived mechanistic descriptors classified as either docking (D), permeation (P), or physicochemical (PC). These are computed using QSAR methods, density functional calculations, ensemble docking and MD simulations in water and in the OM of *P. aeruginosa*. (d) Computational representation of the outer membrane environment (OM) of *P. aeruginosa* detailing the seven sub-regions where MD simulations where the 35 descriptors listed in (c) were computed for each molecule. (e) Principal components third degree decomposition of the molecules following the color code shown in (a).

## 2 Results

Following the protocol outlined in [37], mechanistic descriptors are computed using variety of approaches for a much broader spectrum of molecules. We use traditional chemical/physical property evaluations, density functional theory calculations, and all-atom MD simulations of compounds in water. We refer to these 73 physicochemical (PC) descriptors that depend entirely on the compounds as QSAR, QM, and MD, respectively. In addition, we calculate an additional set of 101 descriptors that are generated based on the interaction of compounds with the bacterium-specific efflux pump and the OM and we call them mechanistic descriptors. Here, to account for influx, we consider descriptors calculated from the all-atom MD simulations of compounds interacting with the OM model [Fig. 1(d)], and, to account for efflux, we consider ensemble docking of compounds targeting specific binding pockets of MexB, the major efflux transporter of *P. aeruginosa*. We refer to them as permeation and docking descriptors, respectively. Our experimental data is obtained by analyzing inhibitory activity of an assembled library of 1260 compounds with antibacterial properties in two mutant derivatives of the wild-type *P. aeruginosa* (PAO1): the PΔ6 strain, which lacks six major efflux pumps (ΔMexAB-oprM, ΔMexCD-oprJ, ΔMexXY, ΔMexJKL, ΔMexEF-oprN, and ΔTriABC), and PΔ6-Pore, which is the hyperporinated version of the PΔ6 strain.

### 2.1 Assembly and properties of the compound library for analyses

For this study, we assembled a unique library of 1260 compounds with antibacterial and efflux inhibitory activities from several different sources [Fig 1(a)]. The library included the two separate compound series developed by Basilea Pharmaceutica [51] (8 compounds) and Rempex Phamaceuticals [52] (255) which were culled from their respective efflux-pump inhibitor (EPI) projects. There were also 92 known antibiotics belonging to various structural classes, including Fluoroquinolone (19) and Beta-lactam (56), were also acquired to be included in this library. Several compound series in the collection were synthesized at Saint Louis University (SLU) as part of on-going EPI and antibacterial projects. The largest source of compounds were a series of EPIs designed to inhibit the AcrAB-TolC pump in *E. coli* including dihydroimidazoline [53] (174), a related series of benzoylimidazolines [54] (14) and a chemical series of 2,7-diaminoquinoline [55] (68). The two sets of antibacterial compounds included a series of 2-aminobenzothiazoles (79) with an unknown antibacterial target and a series of quaternary amine compounds (108) that target a bacterial condensin enzyme [56]. A series of cinnamoyl derivatives [55] (73) which showed both antibacterial and EPI activity were also included in this set.

Several additional series identified in previous screening efforts were also obtained from commercial sources. These included a series of diamino substituted pyrimidines (50) [57], small series of thienylpiperidines and Oxadiazoles (8 compounds each), and a large set of diverse compounds (160) purchased from Enamine. Finally, a set of miscellaneous compounds (161) comprised synthetic intermediates, related analogs that did not belong to one of the aforementioned chemical series and various screening compounds from the NCI collection [58]. We use the 16 chemotype designations to broadly classify the compounds. An alternate classification of the compounds’ 2D structures, using a complete Tanimoto similarity analysis [59], further subdivides the chemotypes shown in Fig. 1(a) yielding a total of 233 subgroups. This high-resolution classification of the chemical structures will become relevant to further analyze permeation predictability of a few key subgroups (see sec. 2.5 and 2.7).

The analyzed 1260 compounds vary in molecular weight (MW) between 156 and 1260 Da, in the total charge between -2 and +5 and have cLogD_7.4_ values between 11 and -11.3. To evaluate the physicochemical space occupied by the library, we carried out the principal component analysis of nine physicochemical properties of the compounds, which included the molecular weight, the number of hydrogen bond donors and acceptors, the total polar surface area (ASA P), clogD_7.4_, the topological surface area, the fraction of sp3 hybridized carbon atoms (Fsp3), the total charge, and the number of rotatable bonds for the analyzed compounds. The first three principal components (PC) [Fig. 1(e)] covered 82.3% of the explained variance. All nine properties almost equally contributed to the compound distribution in the PC1 coordinate, whereas the total charge, the number of hydrogen bond acceptors and the number of rotatable bonds were major contributors in the PC2 [Fig. S1]. Thus, the assembled library is unbiased in respect to one or more features and covers a broad physicochemical space.

### 2.2 Nonlinear relationships among descriptors

The diversity of the chemical space is reflected by the wide range of physicochemical properties of individual compounds, as well as, in their interactions with specific bacterial components such as the OM and the efflux pump. Finding the relevant properties that reliably correlate with a particularly desired behavior or process is challenging: among various descriptors of the compound and molecular descriptors of compound’s interactions with bacterial components, some carry redundant information while others are uninformative. Therefore, reducing the number of descriptors is helpful in developing a robust predictive model. We achieve this by clustering the descriptors and grouping them into subsets that have similar co-variation across the 1260 compounds. Each cluster can be interpreted as a collection of nearly equally informative features, from which one can select a representative covariate to be used to predict an outcome. Such a reduction not only helps manage the complexity of predictive models, but also alleviate the experimental and computational efforts required to characterize each compound.

Traditional correlation coefficients (e.g., Pearson coefficient, *C_ij_*), often implemented by clustering algorithms, quantify the strength of the linear relationships among random variables. Highly correlated variables are expected to belong to the same cluster, while variables with smaller correlation coefficients are placed on different clusters. It is well known that nonlinear transformations of a given variable, while containing the same information, can have a small correlation coefficient. Thus, clustering variables based on the correlation coefficient can have the undesirable property of separating into different clusters, variables that are non-linear transformations from one another. In our evaluation, we observe nonlinear relationships between features, and find that focusing only on linear relationships (e.g., using the standard correlation coefficient to cluster the variables) leads to poor predictive models. To address this problem, we consider rank correlations (*R_ij_*), a generalization of the standard correlation, that captures both linear and (monotone) nonlinear relationships.

Figure 2 depicts key examples of the relationships that are found in the molecular descriptors computed on the studied compounds. The left panel shows the values of the compounds’ cumulative entropy calculated in two different sub-regions of the OM, the lipid surface heads (*y*-axis) and the glycerol basic relationships between descriptors region (*x*-axis). They show a strong linear correlation captured by a Pearson coefficient and rank correlation values close to unity. In contrast, the central panel shows that the relationship between the physicochemical descriptors associated to the rotational constant in the *z* coordinate (*y*-axis) and the isotropic polarization (*x*-axis), is monotonically decreasing. This pair of variables is characterized by a Pearson coefficient of *−*0.4787, which does not quantify the strong non-linear dependency shown. On the other hand, the rank correlation captures better the decreasing monotonic relationship shown by these two physicochemical descriptors with a coefficient value of *−*0.8851. These key changes in the correlation coefficients have greater effects when computing a hierarchical clustering algorithm of the full set of descriptors leading to the identification of 29 clusters using standard correlations, while the rank correlations identify 37 clusters (see SI). In this case, the latter is able to better capture the wide diversity among the different families of descriptors, which has an ultimate key implication when identifying the optimal combination of descriptors (or clusters) that better correlate with the compounds’ desired behavior. Finally, the right panel of Fig. 2 shows the compounds’ molecular weight against their resonantcount, resulting on values close to zero for both measures pointing to uncorrelated variables, which is in agreement with the shown dependency in the plot.

**Figure 2:**
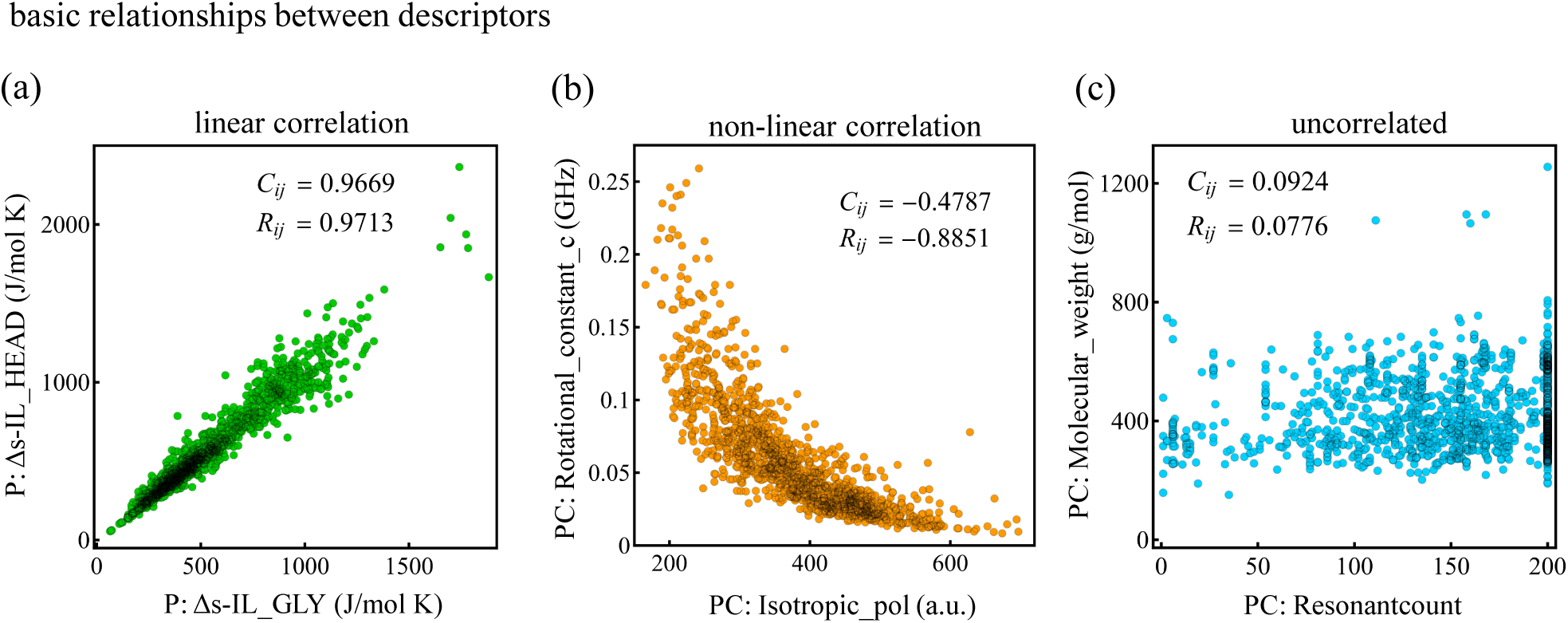
Examples of basic relationships found among the descriptors. Each data point represents a molecule, and it is projected on the two-dimensional space of two descriptors as shown. Some of the most common properties found among the descriptors are: linear correlations (a), non-linear correlations (b), and uncorrelated (c). The numbers on each of the panels are computed by standard correlations (i.e., Pearson coefficient), *C_ij_*, and rank correlations, *R_ij_*. As shown, rank correlations capture better the non-linear relationship shown in the central panel.

### 2.3 Non-trivial relationships among the different classes of descriptors

A hierarchical clustering characterization of the individual families of descriptors reveals two ways in which the grouping of these quantities occurs: first and most simple, descriptors that quantify properties associated with a single attribute gather together, and second, descriptors that are computed in neighboring locations of a specific molecular environment also tend to cluster together. An example of the former is the clustering of size-related intrinsic physicochemical quantities such as the molecular weight and the number of heavy atoms [Figs. S3-S4]. As for the latter, we find that the number of contacts a given compound makes with a specific residue in MexB (docking descriptor) is correlated to that of another residue, if the residues are close to each other within the same MexB monomer [Fig. S5]. Combinations of these two cases are also found. For example, the cumulative entropy associated with a molecule when computed in neighboring sub-regions of the OM (permeation descriptors) are highly correlated among them [Fig. S4]. A detailed quantitative analysis using hierarchical clustering algorithm on these individual families of descriptors is presented in the SI. The natural question that arises is how descriptors, belonging to different families, are correlated with each other, and what is the meaning of these relationships within the context of predicting bacteria permeation and growth inhibition.

Figure 3 shows a visual representation of the individual relationships among all 174 descriptors together with the clusters that are identified by a hierarchical algorithm using the ranked correlation coefficients. The relationship between pairs of descriptors is quantified by means of a dissimilarity matrix (heat map), which is defined as the square-root of the unity minus the square of the (ranked) correlation coefficient associated with the pair, and ordered according to the clustering algorithm (see dendrogram). The colored regions in the dendrogram define the clusters, which are determined by the L-method [60] using the percentage of the variance explained as the critical parameter (see sec. 3.1 in the SI). The procedure identifies 37 clusters in total (blue groups in the dendrogram and white squares in the dissimilarity matrix), 32 of which are comprised by descriptors of a single type, while only 5 clusters are comprised by two or more types. The categorization of the descriptors is illustrated by the following color code: permeation descriptors are magenta, docking descriptors are orange, and for the physicochemical descriptors we further separate them into QSAR in light blue, QM in green, and MD in water in gray.

**Figure 3:**
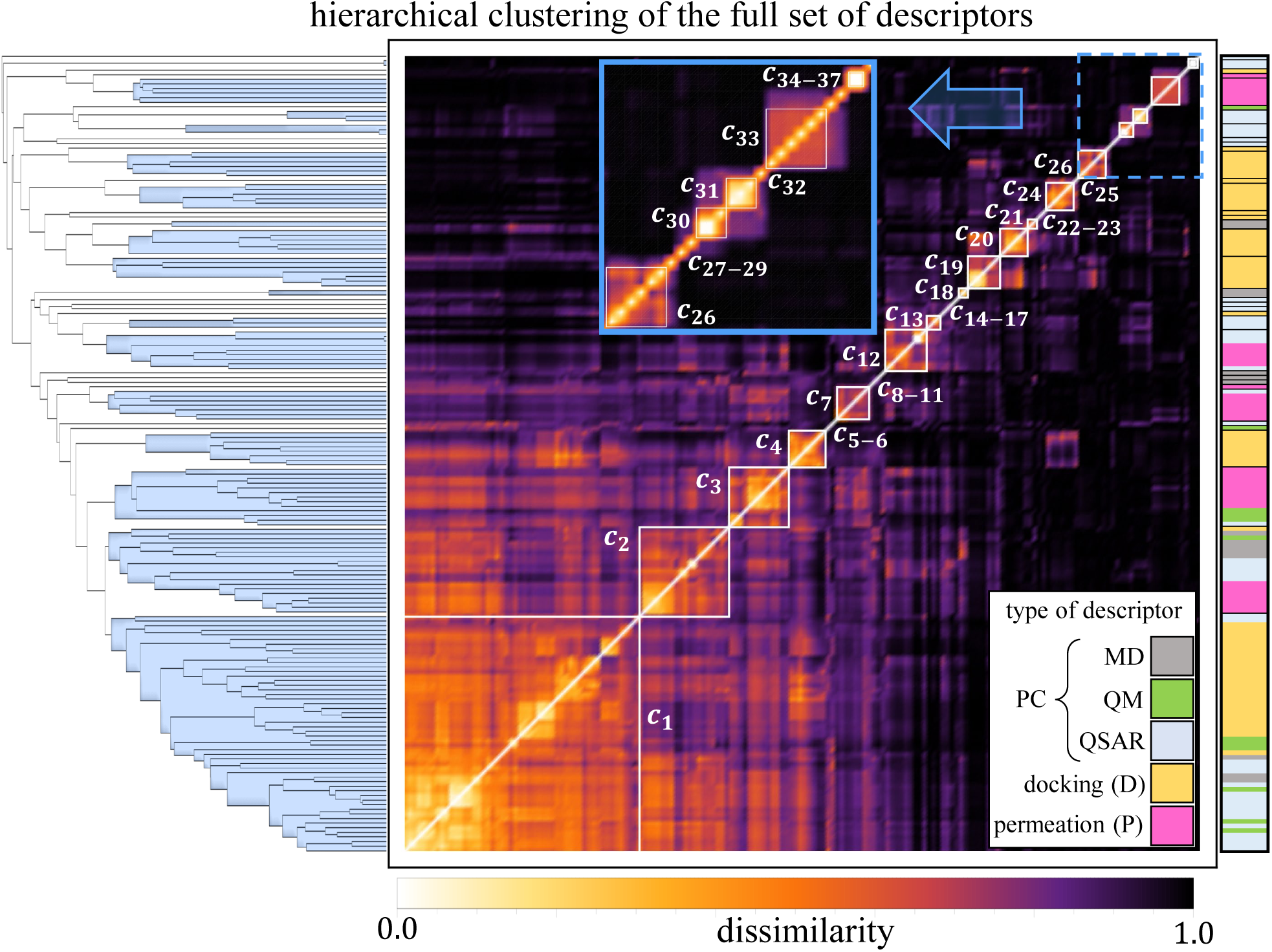
Full data characterization of the 174 molecular descriptors by means of a hierarchical clustering algorithm using their associated rank correlations. The computation yields 37 dissimilar clusters of sizes ranging from single descriptor clusters (e.g., cluster 37) up to a large cluster of 52 descriptors (cluster 1). The dendrogram in the left-hand side depicts the individual as well as cluster level relationships among the descriptors (single line) and clusters (blue groups), respectively. It also permits the visualization of the cut defining the number of clusters, which was determined by the L-method (see sec. 3.1 in the SI). The heat map further highlights the different clusters as well as the relationships between themselves and between individual descriptors via a dissimilarity computation of their associated rank correlations. The type of descriptor is defined in the righthand side by the color code shown in the legend.

Among the single-type clusters found in the full set, there are similar grouping patterns than when performing hierarchical clustering on a single family of descriptors only [Figs. S3-S5], which is expected given the wide spectrum of properties analyzed. However, we also find interesting differences. For example, a single-descriptor cluster in the single-family analysis (e.g., number of donors) becomes part of a larger cluster with descriptors belonging to a different type (e.g., number of hydrogen bonds in OM sub-regions). This provides helpful information to identify redundancy in the information carried by the data, which is desired in order to improve the performance of prediction models. Another interesting finding is that some large clusters formed when grouping single-family descriptors (e.g., permeation only), break when the full set of descriptors is considered. This is also helpful in order to identify outliers with a predictability power difficult to identify when they belong to a larger cluster. This is the case of cluster *c*_8_, which contains one permeation descriptor (HB-MEM-INTER) that is later found to have a great predictability potential. This descriptor, when the clustering is carried out within the single family of permeation descriptors, is part of a larger cluster of other hydrogen bonds-related properties, that together hold a weaker predictability potential. Clusters *c*_33_ and *c*_34_, both of them quantifying the lateral diffusion in sub-regions of the OM, is another example of this type of advantage of using the full set of molecular properties. They comprise a single cluster in the single-family analysis. According to our prediction analysis (presented in the next section), *c*_34_ has a higher potential of becoming a predictor than *c*_33_, and hence, the resultant separation of these clusters in the full set analysis is critical.

When analyzing the full set of descriptors, the largest cluster (*c*_1_) comprises 52 molecular quantities of all types except for permeation. The features grouped in this cluster are mostly related to intrinsic physicochemical properties of the molecules such as size (e.g., volume), graph topology (e.g., Szeged index), polarization (e.g., refractivity), and energy (e.g., thermal energy) of the molecules, together with docking information quantifying the binding energy at both of the studied binding sites of MexB (AP and DP). Interestingly, some additional docking descriptors quantifying the number of contacts between the molecule and residues in DP also comprise *c*_1_. This is explained by a found statistical proximity between these docking descriptors in the DP with the binding energy in both AP and DP [Fig. S5].

Among the clusters with more than one type of descriptors, *c*_2_ is the only one that gathers properties from all types. These include the highly correlated entropy values found in the different subregions of the OM, descriptors quantifying flexibility (rotatable bonds), topology (chain and aliphatic atoms/bonds, rotational constant), and dynamical properties (MD fluctuations and minimal projection area) of the molecule. These properties are found to be correlated also with the number of contacts the molecule makes with residue Thr130 (Threonine) in the deep pocket of MexB. The knowledge of these non-trivial correlations is helpful when determining the predictability power of the cluster, and opens the gate to examine the extent of these relationships in larger families of compounds and the implication when analyzing particular interactions.

### 2.4 Identification of descriptors that predict permeation

The nonlinear relationships found among descriptors, together with the inability of traditional principal component analysis methods to distinguish between weak and strong permeators (see SI), lead us to design a framework that, accounting for these nonlinearities, determines the likelihood that a compound can be classified as be a good or a bad permeator. We aim at identifying a minimal set of molecular descriptors that better correlate with the ability of the compound to permeate/inhibit the pathogen. As many of these properties have strong linear and non-linear correlations among each other, our hierarchical clustering analysis that implements the ranked correlations serves to identify similarities in the descriptor space and hence becomes a good starting point to search for the minimal set of predictors. While descriptors that belong to different clusters are weakly correlated with each other, the information that they carry about the molecule is not redundant and it could point (from different angles) to its ability to permeate and inhibit the bacteria’s growth. Similarly, descriptors within the same cluster carry correlated information about the molecule and not all of these values may be needed.

The target class (i.e., the measure of classification) in these calculations is determined by ratios of the IC_50_ values associated with each compound and extracted from inhibitory activity in two mutant derivatives of *P. aeruginosa* PAO1 (see Fig. 1(b)). For permeation we use the IC_50_ ratio PΔ6-Pore/PΔ6, which highlights the role of the OM. If the ratio tends to 1, the concentration of drug needed to inhibit 50% of the bacterial growth for both derivatives is very similar. This means that the OM barrier makes little to no difference in the action of such drug. Hence, a molecule with such ratio is classified as a strong permeator. Conversely, if the ratio tends to 0, the concentration of drug needed to inhibit 50% of the growth of the PΔ6 derivative is much greater than that needed to inhibit the hyperporinated PΔ6-Pore derivative. Hence, compounds with such ratios are classified as weak permeators. Our calculations show that using a threshold ratio of 0.5 to distinguish between the permeation classes optimizes the classification when compared to other threshold choices (see Fig. S11 in the SI), turning this analysis into a binary classification problem. In short, the target classes are defined as: strong permeators (i.e., class 1) having an IC_50_ ratio greater or equal to 0.5, and weak permeators (i.e., class 0) with an IC_50_ ratio smaller than 0.5. The total number of molecules with measurable inhibitory activity in these mutant pathogens is 602.

Figure 4(a) describes the algorithm designed to reduce the number of descriptors in order to identify an optimal set that are best associated with the molecule’s ability to permeate the bacterial OM. We start by dividing the set of 602 compounds into a large group of 482 compounds and a small group of 120. The large group is used to train/validate on a subset of *x* descriptors a nonlinear classifier and quantify the importance of each descriptor of the subset. The small group is used to test the efficacy of the trained/validated model at predicting the respective target class. The value of *x* is determined by the number of clusters considered in the calculation, and the subset of descriptors is comprised by one descriptor per cluster randomly selected. We start with the complete set of 37 clusters (i.e., *x* = 37 initially). From this set, 200 subsets of *x* descriptors each, are randomly assembled. For each subset, a random forest (i.e., bagged ensemble) classifier [61] comprised by *N_e_*= 1001 estimators (i.e., 1001 classification trees) that use *x*^1*/*2^ descriptors for each estimator assigned randomly with equal weight is implemented using the scikit-learn package in python [62]. Using the properties of the compounds in the training portion, each estimator determines which permeation class is more appropriate for each compound of the testing portion. The dominant class i.e., the one that is assigned by the majority of estimators, becomes the class prediction of the compound in question. The classification algorithm is trained over the target class of 95% of compounds in the large group of compounds, and the remaining 5% is used for validation, which helps to control for over-fitting. For each subset of *x* descriptors, we carry out 50 classification runs over random 95:5 training/validation splits (see dotted circle in Fig. 4(a)). Hence, considering all 200 subsets of *x* descriptors, there are 10,000 classification runs that are carried out for each value of *x*. On each of these runs we measure how good each of the *x* descriptors is at reducing the entropy (i.e., uncertainty) of the classification problem evaluated over the training set. Thus, aggregating the measures of different subsets of *x* randomly selected descriptors, provides a measure of relevance at the level of the cluster. This measure is then used to rank the clusters accordingly. Finally, the number of clusters is reduced by eliminating the lowest scoring one and the cycle is restarted for the reduced set of clusters (i.e., *x → x −* 1). In addition to this process, for each of the classification runs, the fitted model is tested on the small group of 120 compounds (orange arrow in Fig. 4(a)), where we compute the standard confusion matrix and its associated evaluation metrics of accuracy, precision, recall, specificity, and F1. In this way we keep track of how better or worse the model performs for the different combinations of *x* descriptors, as well as, when the number of clusters decreases. An example of the testing portion of the algorithm is illustrated in Fig. 4(b) for the evaluation metric of prediction accuracy (see Fig. S6 in the SI for additional evaluation metrics). For a given value of *x*, it is shown how the random combination of descriptors coming from different clusters perform (orange circles), and how this metric is affected by reducing the number of clusters. It is also shown how the model performance compares with that of a baseline model consisting of simply running the classification algorithm in the full set of 174 descriptors. As illustrated, we find that many combinations of descriptors outperform the baseline model for values of *x* greater than 3. The maximum in the prediction accuracy is found for *x* = 9 clusters for some combinations of descriptors as noted in Fig. 4(c), where the optimal combination of descriptors found is listed (additional details of this calculation are given in Fig. S7). The full ranking of clusters is shown in Table S1 in the SI, where we also show additional details of the model performance for alternative selection of descriptors of these nine clusters [Fig. S8]. Indeed, if we start with a different arrangement of compounds in the large and the small groups, we find changes in the combination of descriptors that maximize the accuracy. Interestingly, the top-9 clusters remain unchanged. The relevance of these nine clusters is preserved even when the large and small groups are randomly scrambled after each iteration (see Table S2). Hence, the information provided by these clusters is crucial at determining the ability of a molecule to permeate and inhibit the pathogen.

**Figure 4:**
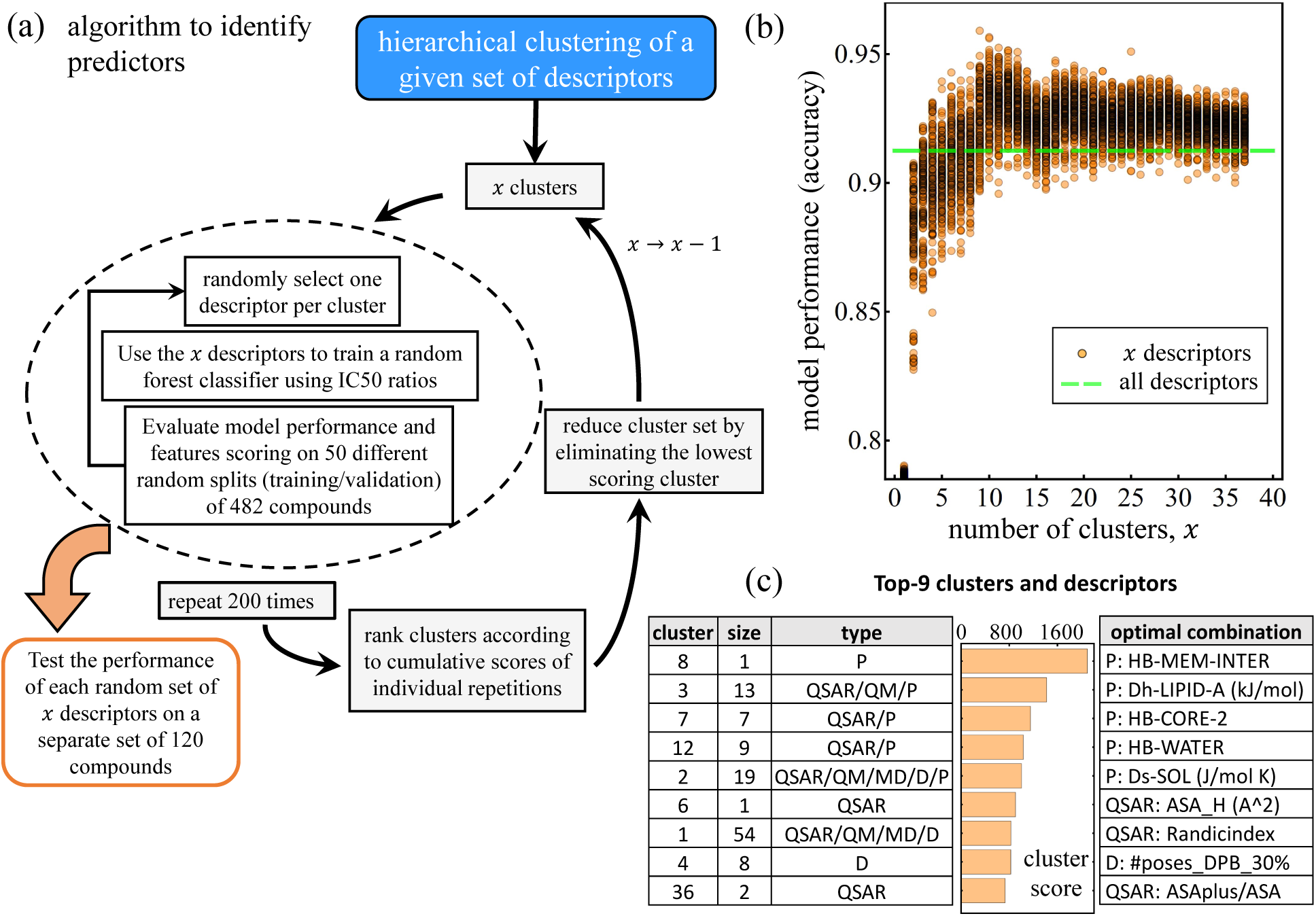
Our model of predictors identification. (a) A hierarchical clustering algorithm is used to select different combinations of *x* descriptors. A random forest classifier is trained on the *x* descriptors alongside with IC_50_ ratios, and the descriptors performance are scored accordingly. Over the course of several random selections of *x* descriptors, the aggregated *x* scores are used to rank the clusters according to predictability. The lowest ranked cluster is eliminated and the value of *x* is reduced. In parallel, for each classification run, the fitted model is tested in a separate set of compounds and the evaluation metrics are stored. (b) Model performance accuracy for each cycle of the model. Individual circles represent the average accuracy score of a single random combination of *x* descriptors using a random forest classifier over 50 random training/validation splits. The dashed green line represents the average accuracy score for a random forest classifier using the full set of 174 descriptors. (c) Top-9 clusters ranked according to their testing performance. The table in the left panel distinguishes the cluster number, its size (number of descriptors comprising the cluster), and type of descriptors they contain. The central panel is the aggregated cluster score where all values add to 10^4^, which is the total number of runs for a particular value of *x*. The right panel lists the top-9 optimal descriptors that produce a testing accuracy of 96.2%.

Fig. 4(c) also shows the ranking of these 9 clusters by their cumulative scores (the sum of all values is 10^4^), and the optimal combination of descriptors that yields a maximum prediction accuracy evaluated in the testing set equal to 96.2%. As shown, within the top predictors, the permeation descriptors associated with the interaction between the compound and the OM of *P. aeruginosa* in the external environment, in the lipid A, and the LPS core 2 sub-regions of the OM, along with the cumulative entropy and number of hydrogen bonds in the water-membrane interface at the inner leaflet of the OM, score the highest. Also, the physicochemical descriptors quantifying the hydrophobic surface area, the ratio between the solvent accessible surface area of all atoms with positive partial charge and the total water accessible surface area (ASAplus/ASA), and the number of docking poses in the DP of MexB complete the list of nine predictors. As mentioned above, the optimal combination of descriptors tends to change with the testing sample [Table S3]. However, we note that this particular combination performs, on average, within 2.3% of the maximum score found for different random testing samples [Figs. S9-S10]. This is encouraging since this particular combination can be generalized across different random testing samples with a modest cost in the performance. Hence, the identification of these markers opens the gate for a deeper study of these key properties, which could guide the design of novel antimicrobials.

### 2.5 Relationship between permeation predictability and chemical structure of compounds

We analyze the robustness of our statistical model and classify the different active compounds in terms of their predictability. We carry out 5000 additional calculations on 100 random testing samples of 120 compounds (50 computations per sample where the training/validation group is scrambled at each computation) employing the classification algorithm over the identified molecular predictors (Fig. 4(c))). The aggregated results of the prediction for each batch of compounds were analyzed and used to classify the molecules in one out of three sets, as depicted in Fig. 5(a). Compounds that were predicted correctly every time, are colored green (set G) and the bar is above the *x*-axis. This is regardless of the compound being a strong or weak permeator, here we are only examining the truthfulness of the prediction. Compounds that are always predicted incorrectly, are colored red (set R) and the bar is below the *x*-axis. Finally, compounds that for some simulations runs were predicted correctly and for some other were predicted incorrectly, are colored blue (set B) and portions of the bar are both above and below the *x*-axis. Interestingly, the largest fraction of the compounds (83.5% or 503 compounds) is predicted correctly every time pointing to a consistent relationship between the predictors’ values and the permeation classes. This regularity is encouraging and talks about the existence of clear trends in the computational data that are strongly linked to the experiments and their consistency across a wide diversity of compounds. We devote the next section to unfold these trends for the most predictive descriptors. On the other hand, sets R and B, though much smaller in size, point to the limitations that a one-rule-fits-all approach have. We find that 9.9% of the compounds (60) were predicted incorrectly every time, pointing to a strong but incorrect signal from the computational data, while the remaining 39 compounds (6.4%) are found to give a mix prediction, which hints to a noisy and hence weak signal. Here we analyze these different sets via our model detailed output and examining the chemical structure of the compounds.

**Figure 5:**
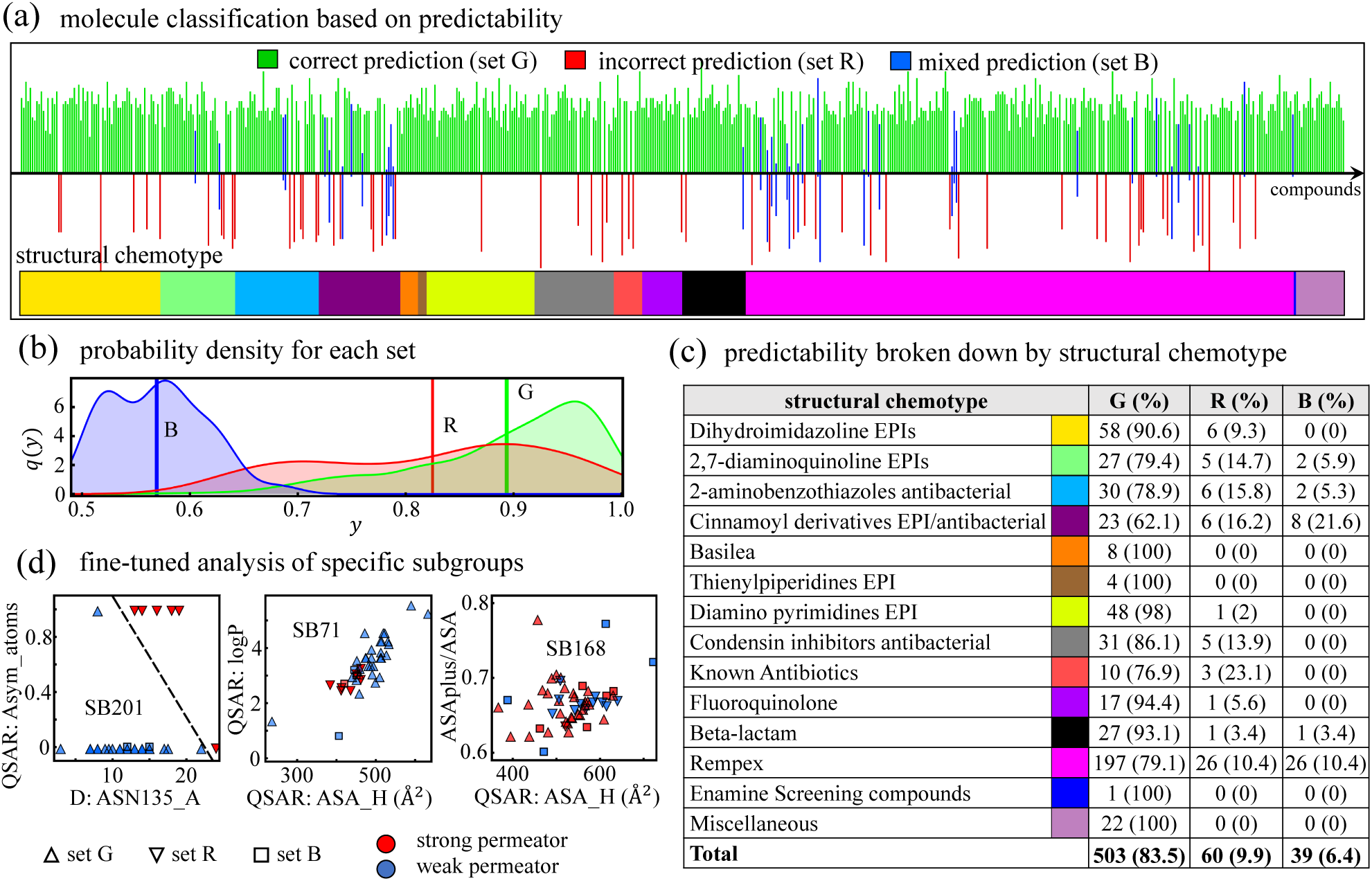
Model prediction analysis. (a) Classification of compounds according to their predictability by our model. 100 random samples of 120 compounds each were tested on the remaining of the data. Compounds that were correctly predicted at each model realization are represented by a green bar pointing above the *x*-axis (set G). Compounds that were incorrectly predicted in every run are represented by a red bar pointing below the *x*-axis (set R). Compounds that in some runs were correctly predicted and in some other, incorrectly predicted, are represented by blue bars pointing both ways (set B). The color bar in the bottom indicates the structural chemotype a given compound belongs to as defined in Fig. 1. (b) Probability density *q*(*y*) as a function of the probability value *y* associated with each category of descriptors (G, R, and B) for the dominant target class, i.e., *y* = max*{p_s_, p_w_}*, where *p_s_* and *p_w_* are the probabilities of being a strong or a weak permeator, respectively. Vertical lines indicate the average probability 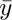 for each case. (c) Number of compounds and percentage of each set (G, R, and B as defined in (a)) for each structural chemotype. (d) Analysis of three selected subgroups according to a complete Tanimoto similarity analysis that contain a relevant amount of compounds from the sets R (inverted triangles) and B (squares). Each panel shows the specific subgroups (SB201, SB71, and SB168) in the space of two descriptors identified by our model (Fig. 4(a)) and compared to their respective experimental class: strong permeator (red) and weak permeator (blue). Dashed line in the left panel is produced by a support vector machine classification algorithm.

To understand the nature of these results, we first analyze the probability densities associated with the identified sets of compounds. We define *y* as the dominant classification probability associated with each compound using the contributions of every estimator for all model realizations. For example, if for a given compound, out of the *N_e_* estimators, *n*_0_ of them choose class 0 while the remaining *n*_1_ = *N_e_ − n*_0_ class 1, we can define the probabilities for each class as *p*_0_ = *n*_0_*/N_e_* and *p*_1_ = *n*_1_*/N_e_*. The dominant probability *y* of the compound is therefore defined as *y* = max*{p*_0_*, p*_1_*}*. The probability density associated with each predictability set, *q*(*y*), is illustrated in Fig. 5(b). Compounds of the set B are characterized by having weak probability values with an average of 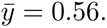 This result is very close to the maximum uncertainty limit of 0.5 (i.e., a coin-flip classification) making the prediction highly unreliable, which is consistent with the mix signal shown in Fig. 5(a). Compounds of the set R have greater values of *y*, but with a wider distribution and an average of 0.82. Finally, compounds of the set G hold a consistently higher average probability of 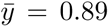 with a median above 0.91. Therefore, having consistently high probability values help rule out compounds from the set B and most of those of the set R.

To explore further these differences, we look at the compounds’ chemical structure and break down the different sets according to the 16 distinct structural chemotypes defined in Figure 1(a). As shown in Figure 5(c), five chemotypes have members in all three predictability sets, five chemotypes have members in two sets, and four chemotypes have members in only the set G. Hence, at this level of analysis, there is not a clear relationship between the structural chemotypes and the predictability sets. A sharper picture can be drawn when we examine the subdivisions of the distict chemotypes by means of a complete Tanimoto similarity analysis. As mentioned in section 1.2, this analysis finds a total of 233 subgroups. Interestingly, nearly 90% of the compounds in sets R and B are concentrated in just 10 Tanimoto subgroups, namely SB71, SB112, SB117, SB118, SB167-SB170, SB201, and SB223. Each of these subgroups is characterized by unique structural features as listed in Table S2 in the supplementary section. We examine the each of these subgroups individually using our model described in Fig. 4(a) but adjusting for the number compounds of each subgroup. In four of these subgroups (SB112, SB170, SB201 and SB223) there is a clear separation between the permeation classes using alternative descriptors to those identified for the full set of active compounds. An example of this finding is illustrated in the left panel of Fig. 5(d) for the subgroup SB201, which belongs to the structural chemotype 3 (i.e., 2-aminobenothiazoles). A combination of the docking descriptor quantifying the number of contacts between the molecule in question and the residue ASN135 in the access monomer in MexB, and the asymmetric atoms allow for a good separation of the permeation classes determined by our experiments in *P. aeruginosa*. This pair of descriptors did not show a wide relevance for the full set of active compounds, but they are found to be key for this specific subgroup of molecules. Our fine-tune analysis of this subgroup correctly classifies the eight compounds of sets R and B that belong to chemotype 3. Similar results are found for subgroups SB112, SB170, and SB223 with different descriptors, as shown in the supplementary section [Fig. S14]. For subgroups SB71 and SB118, the alternative descriptors identified by the model separate better the permeation classes than those of the full set, but some overlap between the classes remains. This is shown in the central panel of Figure 5(d) for the subgroup SB71 (see SB118 in the supplementary section Fig. S14). For the remaining four subgroups (SB117, SB167, SB168, and SB169), which belong to chemotype 14 (i.e., Rempex), a greater overlap between the classes persists pointing to complex nonlinearities between the permeation classes and their descriptors’ values and hence the limitations of using descriptors to find clear trends able to distinguish the molecules according to their permeation class.

Focusing on the set G, which comprises the large majority of active compounds (503 molecules), we find clear trends in the values of the key descriptors that define parameter regions mostly associated to one of the two permation classes [Fig. S12-S13]. For example, strong permeators are found at strong HB interaction values at the surface of the OM (HB-MEM-INTER) together with a strong negative enthalpy in the Lipid-A sub region of the OM. On the other hand, weak permeators fall into the opposite category with weak OM interaction and weak enthalpy [Fig. S13]. In the next section we explore further these general trends and their mechanistic implications in OM permeation.

### 2.6 Descriptor values associated with strong and weak OM permeation

The consistency found in the predictability of the permeation class of the compounds in the set G, makes them a good batch to extract helpful rules that associate specific ranges of descriptor values with a particular target class, i.e., strong or weak OM permeator. To determine the approximate class boundaries and ranges for each individual descriptor, we train a traditional support vector machine (SVM) algorithm [63] for each descriptor of the top-9 clusters (112 descriptors) for the set G of active compounds (503 compounds). Density distribution of descriptor values across the range of selected individual descriptors associated with each permeation class given by their IC_50_ ratio, i.e., strong (red) or weak (blue) permeator, are depicted in Fig. 6. The vertical gray line indicates the binary class boundary identified by SVM. Strong permeators category highly correlates with parameters indicating stronger interactions with different OM subregions, as exemplified in Fig. 6(a). They are able to stabilize a larger number of hydrogen bonds (HB) with different parts of the membrane (e.g. Core-2), while retaining a close water shell during the translocation process in the most hydrophobic regions (e.g. aliphatic tails). Consequently, this contributes with a more favorable enthalpy of interaction in specific parts of the membrane [Fig. 6(b)]. Interestingly, permeation is enhanced by the presence of more rotatable bonds as well as higher entropy, indicative of a more flexible molecular scaffold able to accommodate to the different spatial restrictions along the diffusion pathway. Counterintuitively, larger hydrophobic area does not favor the passage of molecules across the OM, a feature that correlates with the need for localized charges (+2*e* and higher) and stronger dipole moment. Graph-based molecular structure indicators such as Randíc, Harary, Wiener, and Platt indexes are generally higher for strong permeators [Fig. 6(e)]. Finally, for docking descriptors, it is found that strong permeators hold higher number of poses inside the DP in contact with at least *z*% (*z* = 20, 30, 40) of the residues lining the pocket. This yields higher free energy bindings for strong permeators, and a consistently higher number of contacts to key residues inside the DP in MexB, than weak permeators [Fig. 6(f)].

**Figure 6:**
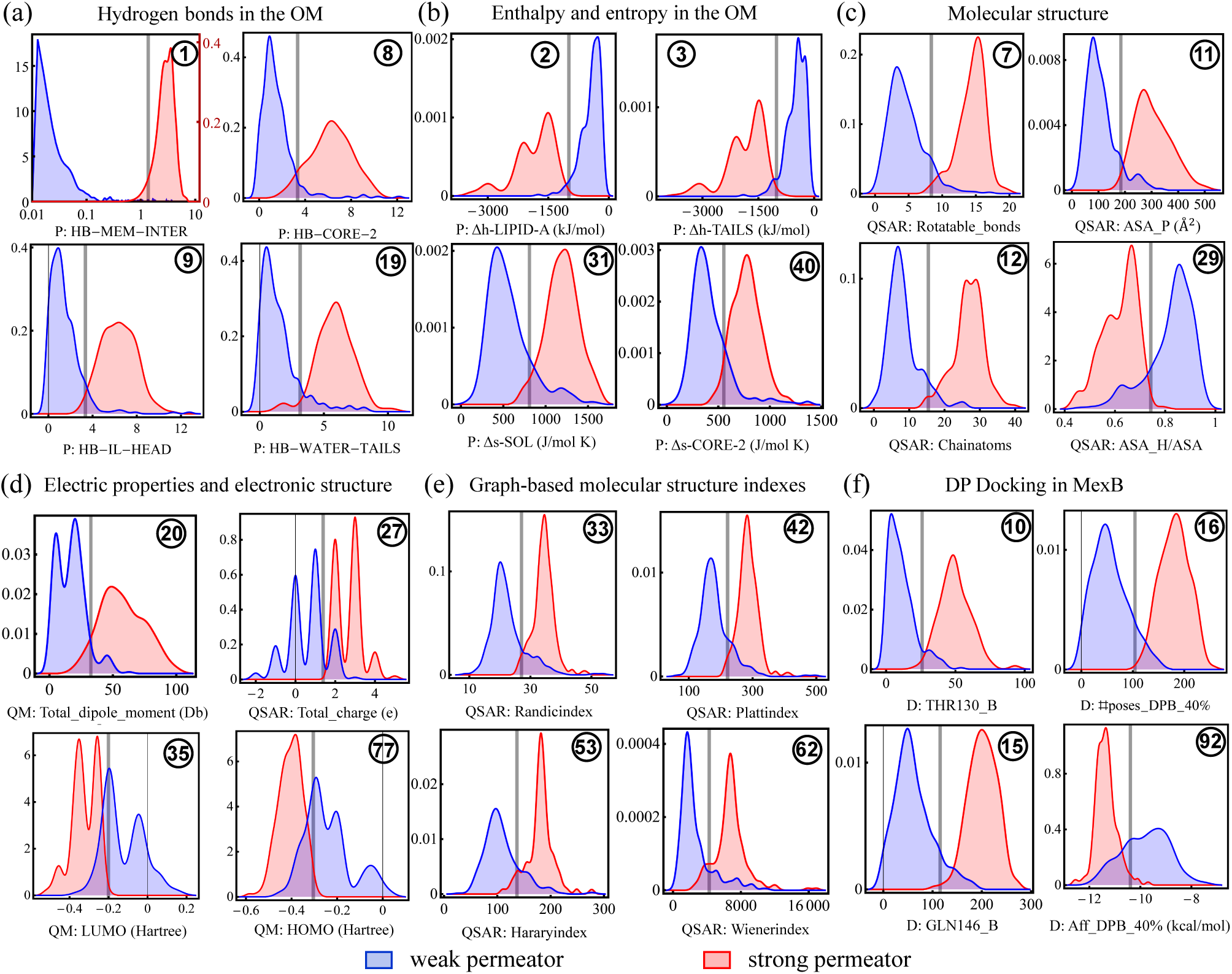
Density distribution values across the range of selected individual descriptors associated with a particular target class given by their IC_50_ ratio, i.e., strong (red) or weak (blue) permeator, for the 506 compounds comprising the predictive group (Fig. 6). The vertical gray line indicates the class threshold estimated by an SVM algorithm. We considered all descriptors from the top 9 clusters from our predictive model (Fig. 4) The descriptors shown hold high predictability scores across general categories (see full list in Tables S4-S5) described as follows: (a) Hydrogen bonds in the OM. Top panel uses two vertical scales and an horizontal logarithmic scale. The red vertical scale corresponds to strong permeators (red). All other panels use a single scale for both categories of compounds. (b) Enthalpy and entropy in the OM, (c) Molecular structure, (d) Electric properties and electronic structure, (e) Graph-based molecular structure indexes, and (f) DP docking in MexB. The circled number in each panel list the ranking according to their single-descriptor predictability scores (Tables S4-S5).

The resultant cutoff values are then tested in the entire set of active compounds (602 compounds) and their associated evaluation metrics are computed for two sets, i.e., set G and the set of all active compounds. These implementations identify simple rules of permeation with very good accuracy in the entire set of active compounds. The circled number in each panel of Fig. 6 refers to the ranking of the descriptor in question according to accuracy for the entire set of active compounds. Tables S4 and S5 list the complete ranking results of the individual descriptors. The resultant one-dimensional ranking is dominated by permeation descriptors. Among those of the top-10, 8 are permeation descriptors. Specifically, the hydrogen bonds and enthalpies computed across different sub-regions of the OM are, overall, the best individual descriptors at determining permeation. Accordingly, and in agreement with our findings at the cluster level, the persistence of hydrogen bonds (time-averaged over 20 ns) between the compound and inner leaflet of the OM at the membrane-water interface (HB-MEM-INTER) is the best single descriptor overall (i.e., best accuracy or *a*_0_), and also the best one at detecting strong permeators (i.e., best positive predictive value or PPV). More than four fifths (0.836) of the active compounds analyzed, including 95.5% of the compounds in set G, that make (time-averaged) 1.36 or more HB with the surface of the outer membrane, are strong permeators.

Among the enthalpy descriptors, and again in agreement with the cluster-level analysis, the enthalpy calculated in the LIPID-A sub-region of the OM ranks the highest in accuracy and in determining strong permeators (i.e., high PPV). More than four fifths of the active compounds (0.815), including 93.7% of the set G, with an enthalpy value in the LIPID-A sub-region smaller than -988.85 kJ/mol, are strong permeators. Combining this information with knowledge of the entropy in the water neighboring the OM (Δs-SOL), increases the fraction of strong permeators correctly identified up to 0.854. Enthalpies associated with other OM sub-regions are next in the ranking of descriptors going from top-3 down to top-6, while the HB in the OM core 2 and heads, are top-7 and 9, respectively. 79.6% and 77.9% of the compounds making 3.34 (time-averaged) HB with the core 2 and heads subregion of the OM, respectively, are strong permeators. The QSAR descriptor quantifying the number of rotational bonds ranks eight and it is the best individual QSAR descriptor overall able to identify 89.2% and 78.06% of the weak and strong permeators, respectively. The best individual docking descriptor is the number of contacts between a compound and the residue THR130 in the DP of MexB. 88.9% of the compounds making less than 26 contacts with this residue are weak permeators. Ranking the compounds according to their negative predictive value (NPV), highlights descriptors that are good at detecting weak permeators. According to this ranking, the molecular orbitals and the number of donors, are the best descriptors at identifying this property. Between 90% and 91% of the active compounds with HOMO (LUMO) levels greater than -0.3045 (-0.2006) Hartree and/or less than four (3.88 on average) donors, are weak permeators.

Next, extending beyond the individual consideration of top-9 clusters (i.e. 1D), we analyze the descriptors for their joint behavior as duets (2D) and triplets of descriptors (3D). Indeed, by carrying out higher order analyzes (i.e., two and more descriptors) it is possible to improve the classification metrics obtained with only one descriptor. For example, combining the permeation descriptor HB-MEM-INTER with the total polar surface area (i.e., QSAR: ASA P(Å^2^)), the classification accuracy increases up to 87.27% while the PPV to 86.68% when tested in the group of all compounds. Further, in a three-descriptor analysis, the highest accuracy score found is 88.4% when combining the enthalpy in the LPS sub-region core 2 (P: Δh-CORE-2), the hydrophobic surface ratio of the compound (QSAR: ASA H/ASA), and the number of contacts that the compound makes with residue PHE615 in the DP of MexB. The PPV and NPV scores are 86.1% and 90.0%, respectively. Certainly, there are many more combinations that produce slightly lower but competitive scores [Fig. S13]. We have listed the most important in the two- and three-descriptor analysis in the Table S6 and S7, respectively, in the SI. According to the analysis in section 2.5, the metric values found for the three-descriptors case lie at the ceiling of the evaluation metric given the behavior of the compounds in the sets R and B. Hence, we do not expect further improvements when going to higher dimensions without breaking down the groups of compounds as we did in the previous section. This is in agreement with the reduction algorithm of the section 2.3, where the random forest determined that nine clusters (*x* = 9) optimizes the classification performance of the entire set, where each classification tree is constructed with the information of three descriptors (i.e., *x*^1*/*2^). The information extracted by this analysis recovers the performance of the nonlinear method (i.e., random forest) and it therefore exhaust the possibilities of better performances with the totality of our data.

### 2.7 Implementation of our statistical model and the permeation rules

As an example of how our analysis can be applied on additional compounds to predict their OM permation, we carry out a testing evaluation on ten new compounds not seen before at any stage of this study. Figures 7(a) and 7(b) detail the structures of these molecules, and show the classification results according to permeation, respectively. The structures and origin of these compounds can be bounded to some of the main 16 chemotypes outlined in Figure 1(a) in the following way: five compounds of chemotype 14 (C2-C6), three of chemotype 1 (C0, C7,C8), one of chemotype 7 (C1), and one of chemotype 9 (C9). Using the most consistent descriptors across several random splits of the training data, the algorithm calculates the probabilities that each compound is either a strong (class 1) or a weak (class 0) OM permeator i.e., *p*_1_ and *p*_0_, respectively. The model calculates consistent prediction probabilities for nine out of the ten compounds. Accordingly, it predicts that four of them are strong permeators (C2-C5) with an average probability of *p*_1_ = 0.94, and five of them are classified as weak permeators (C0,C1,C7-C9) with an average probability of *p*_0_ = 0.84. Experimental results of the IC50 ratios on these compounds in *P. aeruginosa* validates the prediction results for these nine testing compounds. This is illustrated in Fig. 7(b) where we plot the values of *p*_1_ for each compound (black dots) against the target class found experimentally (gray bars). The probabilities calculated for the remaining compound (C6) are *p*_1_ = 0.56 and *p*_0_ = 0.44. These values are very close to those of a random classification (orange line), akin to the values found for compounds of the set B (see Fig. 5(b)). In addition, a structural analysis of this compound indicates that it is akin to subgroup SB167, where the trends between the descriptors and the molecule’s permeation class are not conclusive. The subgroup SB167, which is characterized by having compounds containing an amide derived from 3-aminoquinoline, gathers a large fraction of compounds of the set B. Therefore, we determine that the classification of compound C6 is inconclusive. Our experiments indicate that the compound C6 is a weak permeator.

**Figure 7:**
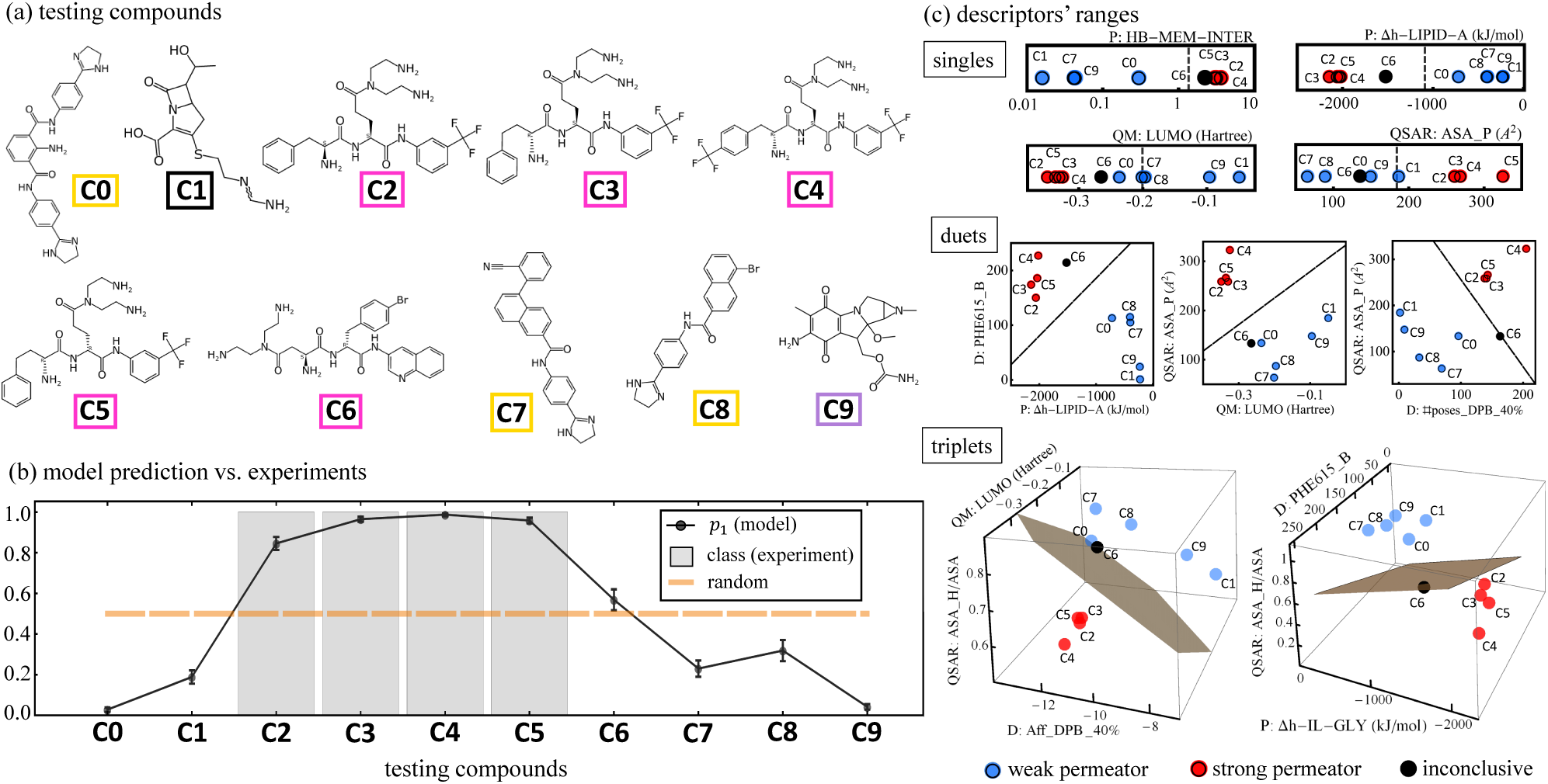
Model testing on additional compounds. (a) Ten compounds labeled C0-C9 structurally classified using the color code defined Figure 1(a). (b) Model prediction associated with the ten compounds (solid black line) against the target class (gray bars) assigned from the IC_50_ ratio measured experimentally in *Pseudomonas aeruginosa*. The prediction quantifies the probability that a given testing compound is a strong permeator, *p*_1_. Error bars are the standard deviation of 100 model runs. Orange line is the maximum uncertainty (i.e., random) classification value of 0.5. The value of *p*_1_ of compound C6 lies very close to this high uncertainty value. (c) Ranges of high-ranked descriptors as singles (top), duets (center), and triplets (bottom) for the testing compounds and classification given by the model. Each panel shows how these compounds’ properties compare to the classification boundary of the training set (dark line or plane).

Figure 7(c) illustrates the values of some key descriptors of the testing compounds as singles (top panel), duets (central panel), and triplets (bottom panel), and how they compare with the permeation boundaries determined in the previous section with the compounds of the training batch. For the high-ranked descriptors (i.e., HB-MEM-INTER and enthalpy in the Lipid A subregion of the OM), most of the testing compounds are in very good agreement with the respective classification boundaries. This further supports our analysis and grants confidence in the applicability to other batches of compounds. Lower-ranked descriptors as singles, such as the LUMO level and polar surface area (ASA P), have an excellent performance as a duet and also as triplets, highlighting an existing complementary relationship among the descriptors. Compound C6 shows an interesting behavior where, for some combination of descriptors, it sides with the strong permeators, while for other, it sides with the weak permeators. This inconclusive outcome with respect to descriptors sheds light on the limitations of an approach based only on these properties to classify some types of compounds. However, having characterized these specific subgroups by their distinguishable structural markers has prevented a possible error in its classification.

## 3 Discussion

Molecular diffusion across bacterial membranes is a very complex process which is largely dependent on the composition of the OM. In particular, Gram-negative organisms have capitalized on very sophisticated mechanisms to ”screen” the passage of molecules from extracellular to intracellular regions. As a consequence, it is extremely difficult to develop new compounds to fight bacterial infections without a proper knowledge of the physical rules governing the overall translocation process. Our results show that small molecule permeation across the OM of *P. aeruginosa* can be predicted with high precision and accuracy based on the abilities of compounds to inhibit growth of cells with the native and hyperporinated OM. These predictions can be made for a library of structurally diverse compounds that likely use different mechanisms to penetrate the OM permeability barrier. However, the robustness of the model is increased by introducing descriptors of passive permeation across the OM model (permeation descriptors). This result provides further support to a recent finding that most antibiotics and nutrients accumulate inside cells by diffusion through the lipid bilayer of the OM.

An accurate calculation of drug permeation is very challenging and requires extensive computing resources as the permeation is related to the exponent of the potential mean force [64]. It becomes impractical when the calculations are required for a large number of compounds such those considered here. Instead, we have generated a large number of molecular and mechanistic descriptors and used machine learning to identify the descriptors that are predictive of compound permeation. Interestingly, we find that descriptors indicative of interactions with different regions of the OM (HB, Δh) are among top ranked predictors for permeation. In fact, a favorable interaction with the membrane can lead to a positive chemical potential in virtue of high compound density, leading to an increase in translocation across the OM [65]. Not surprisingly, the presence of descriptors associated with both the Lipid-A and the core-2 of LPS indicates their importance during passive diffusion. With the exception of the highest ranked descriptor, hydrogen-bonding interactions at the membrane-water interface (HB-MEM-INTER), other highly ranked permeability descriptors can be substituted by QSAR descriptors, resulting in models with slightly lower accuracy. However, this can be beneficial when detailed calculations such MD simulations are not available. This is expected since inherent physicochemical properties of the compounds can be indicative of the chemical space they prefer. In particular, the importance of molecular connectivity (Randic index) and ASA properties for accurate prediction in our analysis indicates that surface exposure to the environment may be critical during the passage of the compounds across the OM. From the mechanistic perspective, the exposure of hydrophobic surfaces (ASA H) can indeed enhance the interaction with the hydrophobic regions of the membrane [66] and is apparently in our calculations a more relevant descriptor for the prediction of permeation than the water-octanol partition coefficients (LogP). We want to highlight however, that QSAR descriptors by themselves are unable to replicate the level of accuracy obtained by using permeation descriptors, highlighting the importance of descriptors generated using all-atom MD simulations of compounds with a realistic Gram-negative OM model.

From a data analysis perspective, our main challenge is to circumnavigate the multidimensional set of descriptors in a way that rationally lessens the computational cost of running a classification algorithm on every single combination of them. The proposed analysis is designed to map, navigate, and reduce this extensive parameter set highlighting the theoretical/computational quantities that better correlate with our experiments. Since the reduction is done at the level of the cluster, it is key to use an optimal clustering technique that accounts for the nonlinearities found in the data. Hierarchical clustering is adequate because it uses correlations as its similarity measure [60] and when applied to a ranked data set (i.e., ranked correlations), it accounts for nonlinear monotonic relationships among the different features [Fig. 2]. In addition, we use a random forest classifier [61] that also identifies and benefits from nonlinear trends found in complex datasets [67–69]. Using a cluster-centered framework alleviates part of the computational cost of testing myriad combinations of parameters and grants a wider perspective on the overall properties that are linked to improvements in their permeation properties. As shown in Fig. S8, although there are performance differences when using different members of a cluster, there are highly correlated descriptors with a performance that are comparable to each other, and the best descriptor is the one that is broadly represented by a general property of the compound (e.g., size) rather than a very specific one (e.g., Wiener index). Even after the implementation of these techniques, the number of possible combinations of descriptors is still very large. A sampling technique was therefore implemented to scan the clusters, rank them according to their predictive capacity, and in parallel, test the different combinations of descriptors of each sample. The effectiveness of this technique at finding a good parameter space for permeation is demonstrated by the more rigorous reverse analysis shown in Figure S7, where alternative combinations of descriptors within the same cluster were identified, and their score compares to that of the combination found during the sampling process. This effectiveness would have not been possible if the clustering technique implemented ignored the nonlinear relationships, since it yields a smaller number of clusters (29 clusters) restricting the parameter space and the descriptors exploration.

Our analysis identified nine key clusters containing the relevant descriptors that maximize the model’s prediction performance, which in turn, allowed us to classify the compounds according to the consistency in their predictability. This classification identified three sets of compounds [Fig. 5]. The largest of them (set G accounting for 83.6% of the compounds) is the most consistent yielding correct predictions in every calculation performed. This gives us confidence in the robustness of the properties captured and in the statistical techniques employed. In reference to the sets R and B, a structural examination using a complete Tanimoto similarity analysis reveals that 10 subgroups contain most of these compounds pointing to a structural connection with their permeation predictability. We find that for some of these subgroups there are subgroup-specific descriptors able to correctly classify the compounds and bypass the prediction difficulty. However, for other subgroups, even the best-ranked descriptors appear to be unable to separate the permeation classes. Though this is a limitation of this approach based on descriptors, this is only found on 4 subgroups of the structural chemotype 14 (Rempex), which is 1 among the 16 structural chemotypes considered in this study. Focusing on the majority set (i.e., set G), we find that it is characterized by projecting strong and weak permeators in well-segregated parameter regions [Fig. S12] allowing us to extract simple empirical rules associated with the descriptor space akin to OM permeation. Using the descriptors of the nine key clusters, we established one-, two- and three-body (i.e., descriptors) rules that better describe the patterns found in all of the active compounds. For example, the one-body analysis highlights the role of the permeation descriptors, especially the hydrogen bonds and the enthalpy computed in several regions of the OM [Table S5-S6]. The patterns found reveal that weak permeators are characterized by having very limited hydrogen bond stabilization with the OM, as well as, having a very weak enthalpy of association [Fig. 6]. The two- and three-body analysis [Figs. S13-S14] revealed a complementary role of the compound’s polar and hydrophobic surface areas that enhance the number of compounds correctly classified [Tables S7-S8]. Docking descriptors, particularly those describing properties of the deep pocket of MexB, are found to be highly correlated with the OM permeation. In fact, there are many examples of three-descriptors sets yielding correct classification scores that are comprised by one permeation, one docking, and one QSAR descriptor [Table S7]. This highlights a complex relationship among these types of descriptors that captures well-rounded properties of both, weak and strong permeation, and hence, it facilitates their correct identification. An application of these uncovered rules on a new batch of compounds demonstrate their predictability power and opens the door to similar data-driven studies in other Gram-negative pathogens. This analysis complements similar efforts at determining the key properties that distinguishes strong and weak permeators [70].

In summary, our work combines experimental, computational, and statistical protocols in order to identify the critical properties that optimally predicts the passage of molecules across the bacterial OM and inhibit growth of Gram-Negative *P. aeruginosa*. The successful approach was able to reduce the spectrum of relevant mechanistic properties in a set of chemically diverse compounds with known antibacterial activity into simple but non-trivial empirical rules for the prediction of strong or weak permeators. We hope that the found relationships can guide additional experimental efforts and accelerate the rational design of new classes of molecules for combating antibiotic resistant strains. Our approach can be expanded for targeting the permeability of molecules to different biological membranes, regardless of their composition or distribution.

## Supplementary Information

### 1 Experimental Methods and Chemical Syntheses

The experimental set-up has been reported before [1, 2]. Briefly, *P. aeruginosa* cells were grown in Luria Bertani Broth (LB) (10 g tryptone, 5 g yeast extract, 5 g NaCl per liter, pH 7.0) at 37° C with shaking. Inhibitory concentration (IC_50_) determination was carried out using the 2-fold broth dilution method. Two independent experiments were carried out. The expression of the Pore was induced at OD_600_ = 0.3 *−* 0.4 by addition of 0.1 mM IPTG. Chemical structures of the assembled library of 1260 compounds and the measured IC_50_ values are available upon request.

#### 1.1 Principal component analysis of our library of compounds

As mentioned in the main paper, the library is comprised of 16 different structural groups coming from different sources, including known antibiotics. A principal components decomposition using nine basic properties properties is shown in Fig. 1(d) of the main paper. Figure S1, shows the contribution of each of these nine properties to the first three principal components. This is calculated from the eigenvectors of the covariance matrix of the standardized data (i.e., after a z-score normalization). The absolute value of the entry of each eigenvector is scaled so that the sum is 100, i.e., a percentage. As shown, the first principal component has a rather similar contribution of each of the properties. On the other hand, the corresponding eigenvalues provide information about the variance held by each degree of the principal components. In Fig. 1(d) of the main paper, we show the percentage held by the first three.

**Figure S1:**
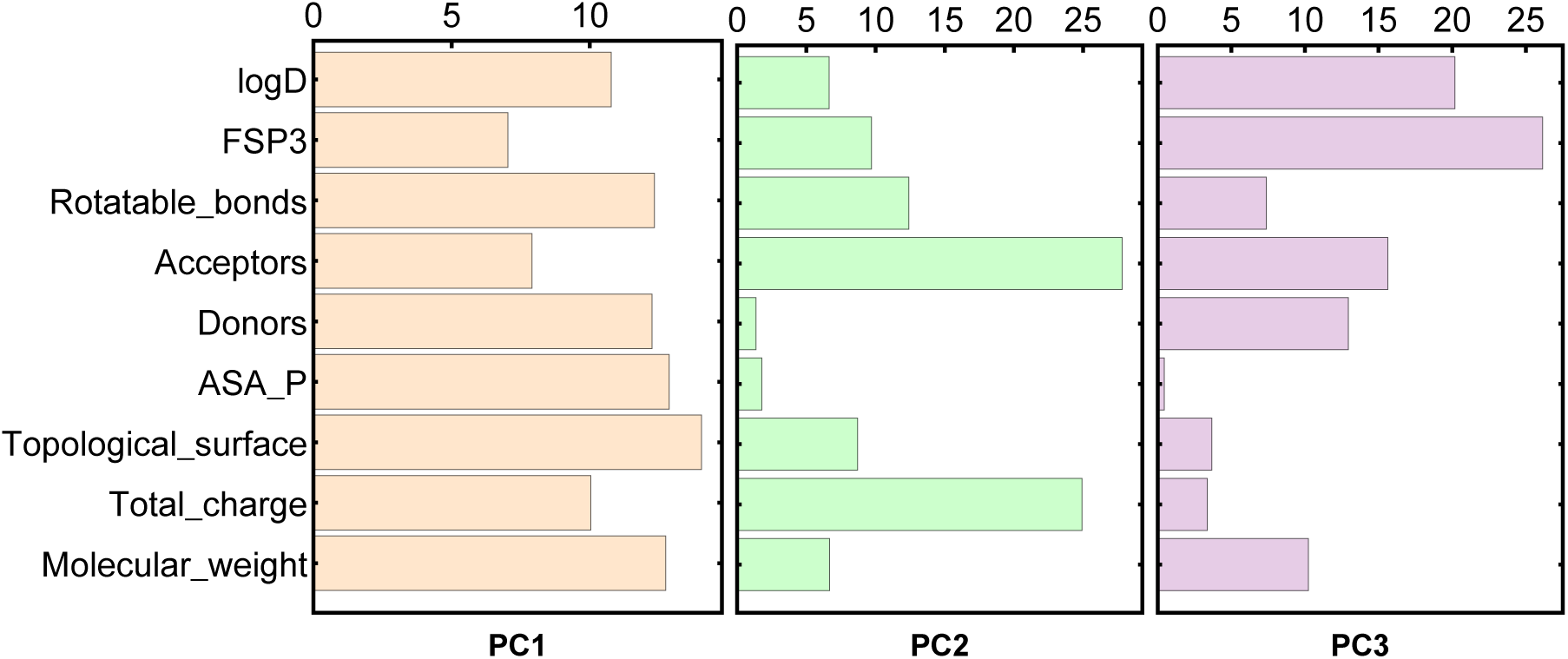
Contribution of each of the nine listed properties to the principal components representation up to third degree for the full set of 1260 compounds used in this study. The contribution is calculated from the eigenvectors of the covariance matrix.

### 2 Computational Setup and Protocols for Computing Molecular Descriptors

#### 2.1 QSAR, QM, and MD calculations

For each compound we considered the protonation/charge state most populated at physiological pH. We used the ChemAxon’s Marvin suite of programs [3] to obtain standard 1-2-3D descriptors used in QSAR studies (e.g., numbers of heavy atoms, rotatable bonds, H-bond donors/acceptors, van der Waals volume and surface, etc. see [1]). The geometry of the major microspecies has been used to perform QM calculations with the Gaussian16 package [4] as previously described [5]. Employing a polarizable continuum model to mimic the effect of water solvent we optimized the ground-state structure and performed full vibrational analysis, obtaining real frequencies in all cases. On the optimized geometry, we performed single-point energy calculations in vacuum to generate the atomic partial charges fitting the molecular electrostatic potential.

Under the constraint of reproducing the electric dipole moment of the molecule, we used the Merz-Kollman scheme [6]. Atomic partial charges were generated through the two-step restrained electrostatic potential method [7] implemented in the AnteChamber package [8]. With this program we derived general Amber force field (GAFF) parameters [9]. QM descriptors associated with the ground-state optimized structure include static polarizabilities, frontier molecular orbital energies, permanent dipole moment, and rotational constants. For each compound, we performed 1-*µ*s-long all-atom MD simulation in explicit water solution (0.1 M KCl) using the Amber18 package as described before [5]. From MD simulations, we obtained structural and dynamic features of the compounds investigated by means of the CPPTRAJ program [10]. The number and population of structural clusters were determined using a hierarchical agglomerative algorithm [11].

#### 2.2 *P. aeruginosa* OM set up for MD

The initial coordinates of the Outer membrane (OM) corresponding to the *P. aeruginosa* Gram-negative bacteria were downloaded from http://dqfnet.ufpe.br/biomat/software.html. The model has been parameterized in line with the GLYCAM force field[18] and parameters are adapted to run in the GROMACS[19] molecular dynamics engine. Briefly, the OM consists of an inner leaflet composed of 1,2-dipalmitoyl-snglycero-3-phosphoethanolamine (DPPE) and an outer leaflet composed of a truncated LPS structure. The membrane is fully solvated using the TIP3P water[20] model and anionic charges in the LPS molecules are counter balanced with CA++ cations. A schematic representation of the model is provided in Fig. 1(e) in the main paper and more details about its parameterization can be found in the original work[21].

Compounds were represented using the Amber force field. First, we optimized the ground-state structure of each compound employing a polarizable continuum model [22] as to mimic the effect of water solvent particularly to avoid formation of strong intramolecular H-bonds. This geometry was confirmed performing a full vibrational analyses, obtaining real frequencies in all cases. On the optimized geometry, we then performed single-point energy calculations in vacuum to generate the atomic partial charges fitting the molecular electrostatic potential. Under the constraint of reproducing the electric dipole moment of the molecule, we used the Merz-Kollman scheme [23] to construct a grid of points around the molecule. Atomic partial charges were then generated through the two-step restrained electrostatic potential method [24] implemented in the AnteChamber package [25]. Using this program, we derived general Amber force field (GAFF) parameter[26], which were transformed into GROMACS input files using the antechamber python parser interface (ACPYPE) tool[27].

##### 2.2.1 MD protocol for computing the permeation descriptors

In order to screen the molecular descriptors corresponding to the permeation along the OM membrane, each drug was placed into seven different molecular environments corresponding to specific regions along the normal of the OM (Fig. 1(e) in the main paper). These regions were explicitly selected in order to cover the influence of both the inner (DPPE) and outer leaflet (LPS) of the OM. Thus, seven independent simulations per drug were necessary in order to recapitulate the influence of the OM into the permeation process. The whole procedure was automated via a series of bash scripts, which iteratively connected the pulling code and energy minimization in GROMACS[19].

All simulations were run with the GROMACS 5.4.1 molecular dynamics engine[19] with a time step of 2 fs. The LINCS algorithm[28] was applied to constrain all bond lengths with a relative geometric tolerance of 10^−4^. In line with its original parameterization, short-range interactions (vdW and Coulomb) were calculated using a cut-off scheme of 0.9 nm, which were evaluated based on a pair-list recalculated every five time steps. Long-range interactions were handled using a reaction field[29] correction with a permittivity dielectric constant of 66. After initial set-up, each system was energy minimized using 3000 steps of conjugated gradient, followed by a thermal equilibration of 1 ns. A harmonic potential of 1000 kJ mol-2, along the Z vector connecting the center of mass (COM) of the drug and the OM of the membrane was applied in order to maintain the relative position of the drug with respect to each of the seven defined regions of the membrane (Fig. 1). During equilibration, bilayers were coupled to 1.0 bar using a Berendsen barostat[30] through a semi-isotropic approach with relaxation time of 1.0 ps. Afterwards, production runs were coupled using a Parrinello barostat[31] algorithm and a constant temperature of 310K was maintained by weak coupling of the solvent and solute separately to a velocity-rescaling[32] scheme with a relaxation time of 1.0 ps. Production simulations were run for 20 ns and trajectories were saved each 20 ps.

A total of 8841 (176 *µ*s) trajectories were analyzed using in-home developed bash scripts, which were directly interconnected to the in-built GROMACS tools. Thus, for each simulation the following molecular descriptors were evaluated (Fig. 1(e) in the main paper): Number of hydrogen bonds between the drug with its first solvation shell (HB-WATER), number of hydrogen bonds between the drug and the surrounding OM environment (HB), lateral mean squared displacement of the Drug (Δxy), Total enthalpic component of interaction between drug and surrounding environment (Δh),and total cummulative entropy of the drug (Δs). All these analysis were carried with the in-built analysis tool set provided in GROMACS.

#### 2.3 Ensemble docking to MexB

Molecular docking calculations were performed using the AutoDock Vina package [12]. The program was used with default settings except for the exhaustiveness parameter which was set to 1024 (default of 8). Protein and ligand input files were prepared with AutoDock Tools [13]. Flexibility of docking partners was considered indirectly by using the ensemble of conformations. In particular, for each compound we used 10 different cluster representatives extracted from MD simulations in explicit water solution, while for MexB, we considered 6 conformations, including available X-ray crystal structures (PDB Ids 2V50, 3W9I, and 3W9J) [14, 15] and MD snapshots extracted from MD simulations [16]. For each docking run, we retained the top 10 docking poses. Following Ref. [17] we performed two sets of guided docking runs into the two major binding pockets of MexB: the access pocket of the access monomer (AP) and the deep binding pocket of the binding monomer (DP). In each case, the docking search was performed within a cubic volume of 40*×*40*×*40 Å^3^ centered in the center of mass of the pocket. The interaction between compounds and MexB was quantified by means of a statistical analysis of all poses, yielding about 60 descriptors. These descriptors include average binding affinities (according to the docking scoring function) as well as the total number of contacts with single residues lining the two pockets (see Table S1).

### 3 Hierarchical Clustering Implementation

#### 3.1 Agglomerative clustering: generalities and applications

This is an unsupervised statistical technique that uses correlations among random variables to form groups (or clusters) of highly correlated quantities, resulting in clusters that are highly dissimilar from one another. This is a bottom-up technique that starts with clusters formed by a single random variable. Then the correlations coefficients among all the pairs are computed and ranked. The pair with the lowest dissimilarity measure is merged together into a cluster of size two. The dissimilarity *D_ij_* is defined as the square-root of one minus the square of the correlation coefficient between the pair *i* and *j*:

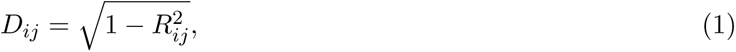

where, *R_ij_* is the correlation coefficient between variables *i* and *j*. Subsequently, all correlation coefficients are computed again treating the cluster of two as a single variable in which the resultant correlation between the pair and another variable is derived as the average of the correlation with each member of the cluster individually. Then, the dissimilarity measures among all groups are ranked and the pair with the lowest one is merged into a larger cluster. This process is repeated until only one cluster remains.

In our analysis we implement the ranked correlations coefficients consisting on replacing the value of the random variable (i.e., molecular descriptor of the compound) for the low-to-high rank of such value within the distribution. For example, for the molecular property of molecular weight, the lightest compound would have a rank of one, the second lightest a rank of two, and so one. We do the same procedure for all descriptors. Then, the ranked correlations are calculated by computing the Pearson correlation coefficient over the list of ranked values.

In order to determine the optimal number of clusters we use the fractional variance explained defined as the ration of the variance between groups (i.e., points residing in different clusters) to the total variance (i.e., all points):

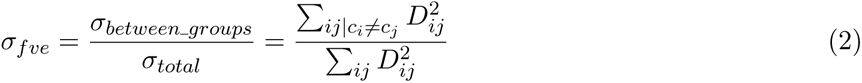

This quantity increases as the number of clusters decreases and then stabilizes, which points to an appropriate number of clusters. At this point variance from within clusters is small enough hinting at a relative closeness among points within clusters and otherwise for points in different clusters. The L method [**?**] is employed to identify the optimal number of clusters *n_c_*. First we create the list of the fractional variance explained *σ_fve_* vs the number of clusters *n*. For each candidate number *n* we find the best straight line fit of all points before and after *n*, and compute the weighted sum of the root mean square error (RMSE) associated to the fits. The value of *n* that minimizes RMSE corresponds to the point in which the variance stopped increasing as a function of the number of clusters. We consider this point as the optimal number of clusters. Figure S2 illustrates the application of this technique to determine the optimal number of clusters for each set of descriptors independently as well as for the combined set.

**Figure S2:**
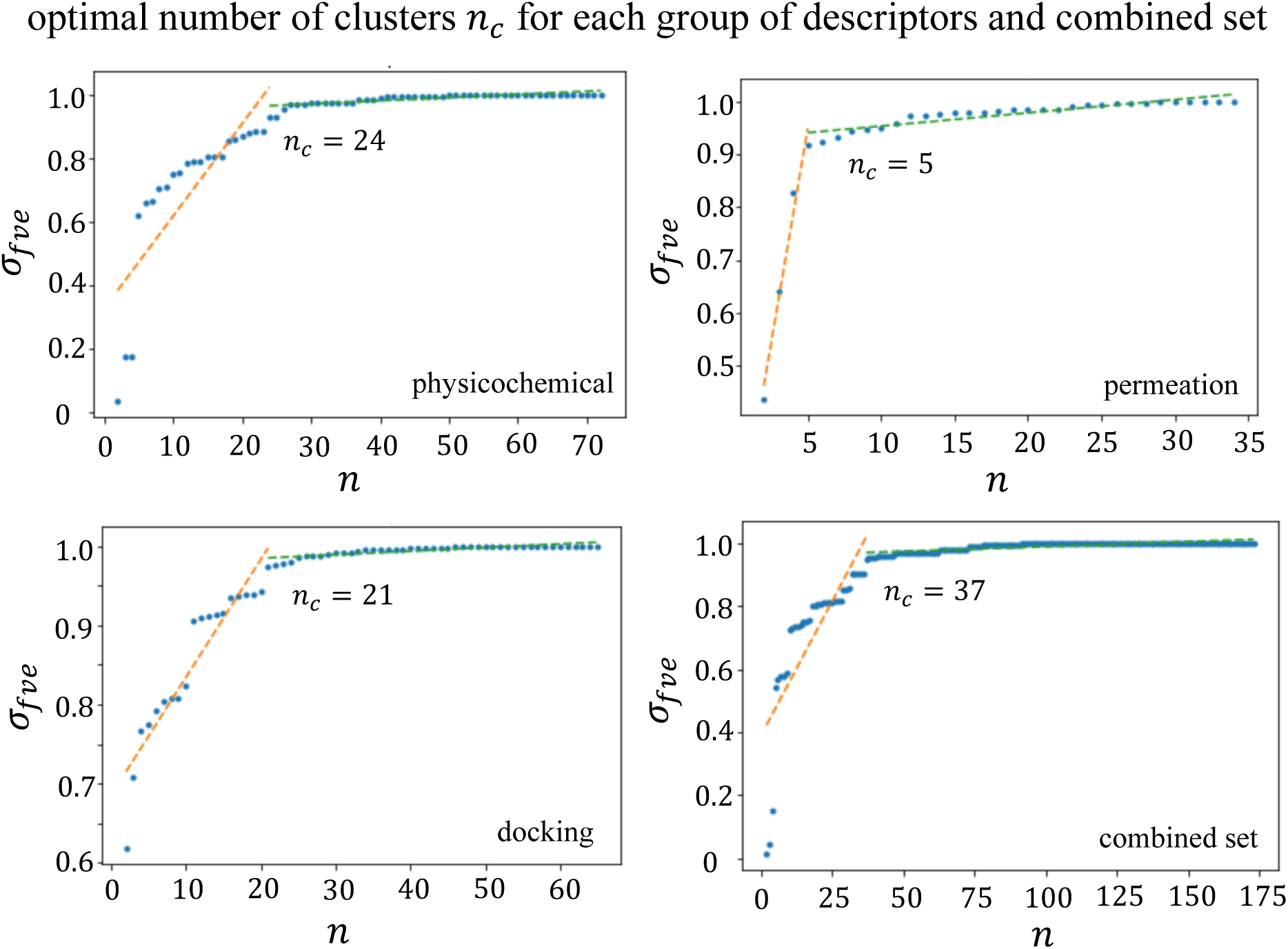
Finding the optimal number of clusters for each set of molecular descriptors and for the combined set following the L method

#### 3.2 Physicochemical and permeation descriptors cluster around the type of observable

We compute non-linear hierarchical clustering to the group of 73 physicochemical descriptors, as well as, to the group of 35 permeation descriptors using the ranked correlations and the full set of compounds (see Fig 1). For both cases we find that the resultant clusters are formed around descriptors that quantify similar observables. This is not surprising for the physicochemical descriptors given that these quantities measure intrinsic properties of the compound (e.g., total charge), or they were computed in simple water environments. Fig S3 illustrates the classification of the 73 physicochemical descriptors in 24 clusters. The largest of them comprises 32 descriptors, which quantify properties akin to the size of the compound (e.g., molecular weight, atom count, and volume), together with positive, negative, and water accessible surface areas. These are mostly QSAR and QM descriptors. Slightly less correlated, but still within the same large cluster we find descriptors computed via MD simulations in water environments. These include the average value of the minimal projection area associated to the configurations explored by the molecule in the MD trajectory, and the average value of the root mean square fluctuation (RMSF) of the atomic positions. On the other hand, the remaining clusters are comprised by between one and five descriptors. These clusters group additional compounds properties such as charge and molecular orbitals, partition coefficients, shape, surface, and aromatic properties (full list given in the SM). Most of these smaller clusters contain only one type of descriptor (either MD, or QSAR or QM), and highly dissimilar among each other as depicted in the heatmap and dendrogram of Fig S3.

**Figure S3:**
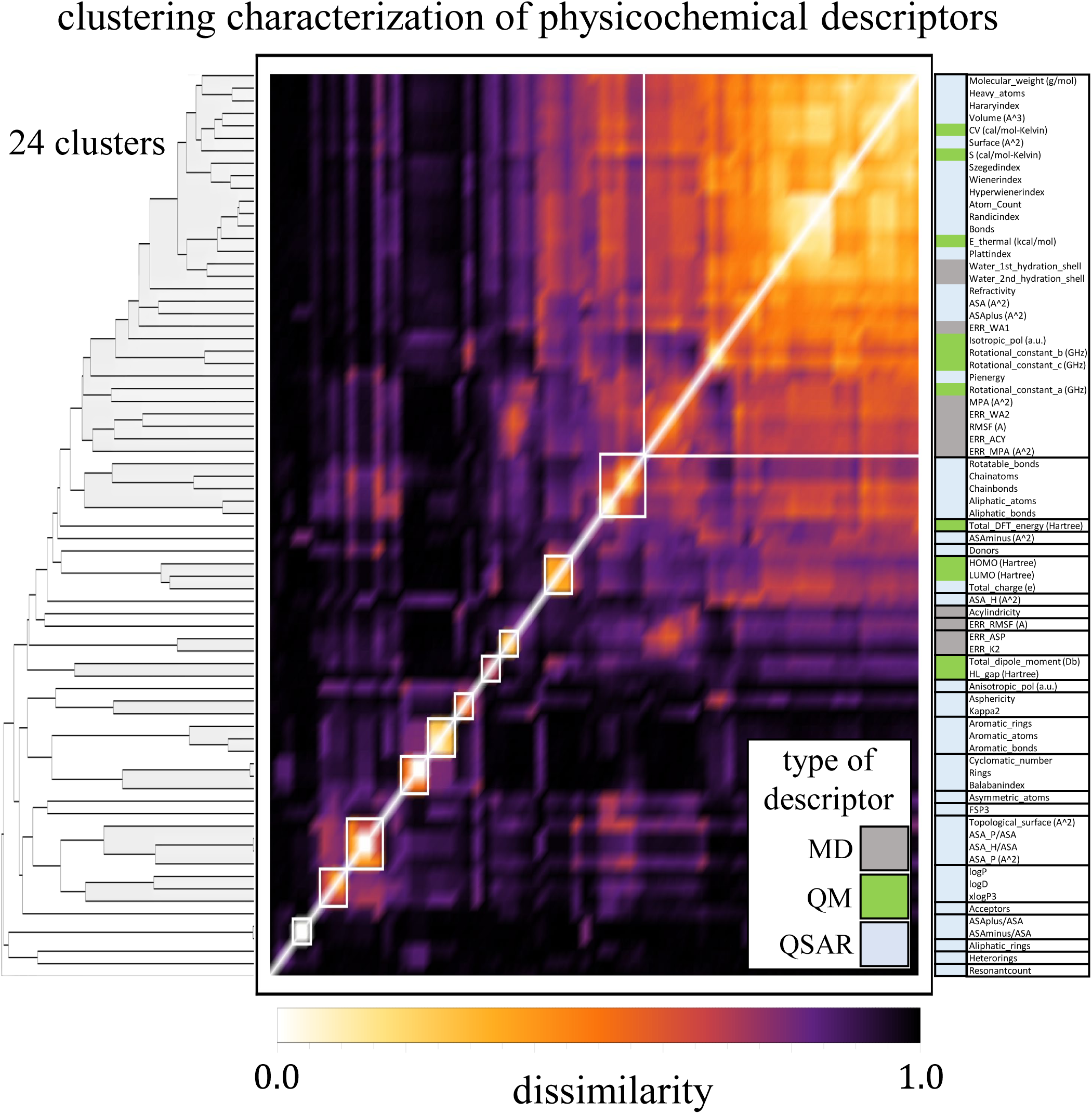
Clustering characterization of physicochemical descriptors. A hierarchical clustering is computed with the ranked correlations of 73 physicochemical descriptors for each of the 1263 compounds derived from quantitative structure-activity relationship (QSAR) modeling, density functional quantum mechanical calculations (QM), and molecular dynamics (MD) simulations in water, as shown in the color bar at the right hand side. The calculation yields 24 clusters arranged by the dendrogram and illustrated by heatmap by means of the dissimilarity defined as 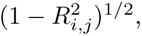 where *R_ij_* is the ranked correlation between descriptors *i* and *j*.

On the other hand, the permeation descriptors were computed through extensive all-atom MD simulations in seven different regions of the simulated OM of P. aeruginosa. These regions are characterized by having contrasting chemical properties and dissimilar compounds interact very differently with each of the region. In spite of this contrast, our clustering analysis groups together descriptors quantifying the same type of observable (e.g., entropy) measured at the different OM regions. This is in contrast with our analysis of the docking descriptors where it was found that the region where these properties were computed defined the cluster these belong to. Permeation descriptors group according to observable rather than the region. The only exception found is the number of hydrogen bonds between the compound and surrounding water molecules in the hydrophobic tails region, as well as, the lipid A sub-region. Our analysis finds that these descriptors are statistically more akin to the interaction energy (i.e., enthalpy Δh) between the compound and the OM than to the hydrogen bonds in the other regions. Moreover, hierarchical relationships among neighboring regions of the OM are found when we look at highly correlated descriptors belonging to the same cluster. The clearest example is the clustering of the entropy values, where we find a hierarchy of correlations among descriptors that resemble almost perfectly the neighboring regions in the OM. The most correlated pair is the entropies associated with the two outer sub-regions (SOL and HEAD), which in turn, is highly correlated to the third outer region of Glycerol. The resultant group is statistically more akin to the hydrophobic tails, forming a group that is more alike to lipid A. Finally, the resultant group is associated to the core regions. The high correlation between an observable measured in neighboring regions of the OM can be found in all clusters.

#### 3.3 Correlations in the number of contacts between compounds and key residues in MexB are primarily associated to residues proximity

A statistical analysis of the docking descriptors using hierarchical clustering and rank correlations results in 21 clusters of different sizes ranging from single descriptor clusters and up to one cluster comprised of 13 descriptors. This is represented in the dendrogram of Fig S5. As reasonably found, descriptors associated to DPB tend to cluster together and are highly dissimilar statistically from those associated to APA, which also clustered together among themselves. These two highly dissimilar groups break further into clusters of descriptors, some of which quantify the number of contacts to specific residues, while other the access and binding energies. A direct examination of the clusters containing the descriptors that quantify the contacts to key residues and the structure of the APA and DPB protomers, clustered residues tend to be in close proximity to each other within the protomer. The residues contained in the clusters are not characterized by similar physicochemical properties, but interestingly, the clusters also identify different sub regions of both of the pockets. For example, those in the DPB are close to the hydrophobic trap and appear to line a region in which compounds can fit well. In fact, the group of residues in the DPB are all hydrophobic, which is also consistent with what is expected since hydrophobicity is a common feature of good substrates/inhibitors of the transporter [36, 37]. On the other hand, those of the APA represent the sub-region just before entrance to the DPB, close to the G-loop separating the two pockets. In a like manner, we also find strong correlations among descriptors quantifying the average binding affinity (as predicted by docking) in the access and distal pockets however they are within the same cluster.

**Figure S4:**
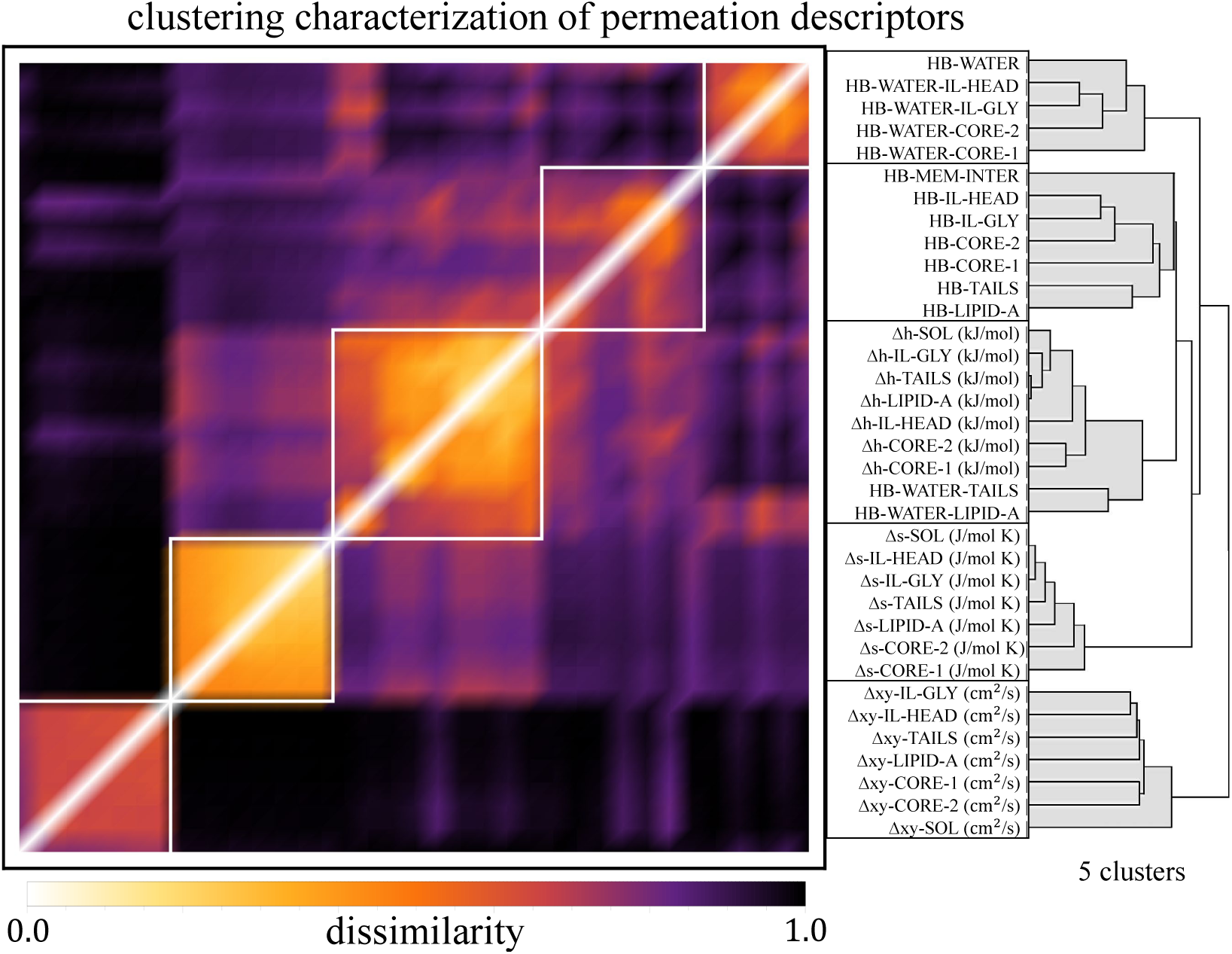
Hierarchical clustering characterization of the permeation descriptors revealing 5 dissimilar clusters grouped, for the most part, by physical quantity.

**Figure S5:**
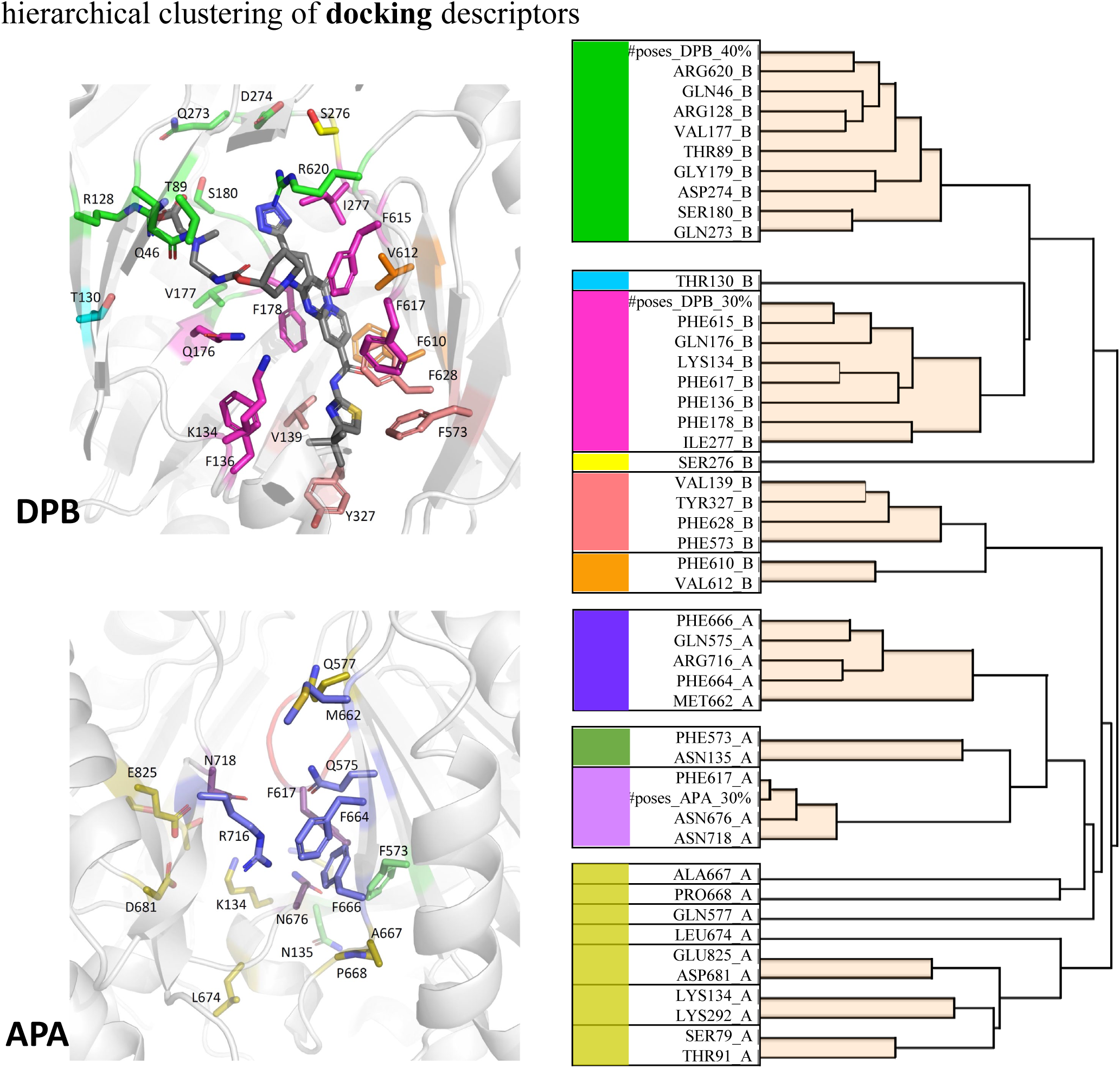
Hierarchical clustering of docking descriptors showing a subset of those associated to contacts to residues in MexB. Our statistical analysis yields 21 clusters in total. For illustration purposes, we present 16 of them that group the number of contacts between the compound and specific residues in the DP T (upper left) and AP L (bottom left) pockets of MexB. There is a clear separation between the descriptors associated to residues in the DP T and those in the AP L, showing stronger similarities in the former, as shown the dendrogram (right panel). These residues depict clear correlations between proximity and cluster membership.

### 4 Full ranking of clusters according to predictability of OM permeation using the full set of descriptors

Table S1 lists all of the clusters, their low-to-high ranking according to prediction of permeability, and the descriptors that comprise each of the clusters as illustrated in the Figure 3 of the main paper. Complementary of this ranking, we repeat the process but we now randomly produce a new training and testing sample after each model iteration. The ranking up to top-21 is shown in Table S2. As shown, the top-9 clusters remains stable and their relevance is not dependent in the specifics of the testing sample employed for the data analysis.

### 5 Our reduction algorithm and its robustness across combinations of descriptors

#### 5.1 Evaluation metrics: definitions

The model evaluation metrics computed in this work are based on combinations of the output from the traditional confusion matrix, which compares the fruitfulness of the prediction (i.e., true or false), with the binary classification (class 1 or class 0) of the real data. Hence, each prediction outcome can be classified as either true positive (*TP*, or class 1 correctly identified), true negative (*TN*, or class 0 correctly identified), false positive (*FP*, or a real class 0 identified as class 1), or false negative (*FN*, or real class 1 identified as class 0). The accuracy (*a*_0_) is the ratio of correct predictions to all predictions, i.e., the fraction of correct predictions:

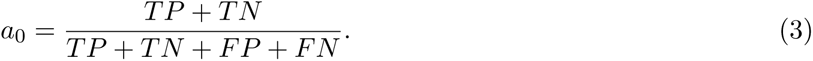

Similarly, the classification error (*e*_0_) is defined as the opposite to the accuracy, which is the ratio of the incorrect predictions to all predictions, i.e., the fraction of incorrect predictions:

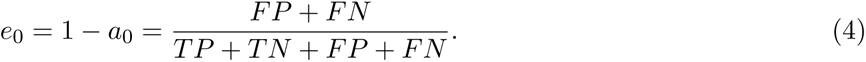

In addition, metrics targeted to reduce specific error outcomes provide different angles of a given classification model. For example, recall (*r*_0_) calculates the fraction of all positive instances that are correctly identified, and hence it is also known as the true positive rate (TPR). Its purpose is to minimize the false negative outputs:

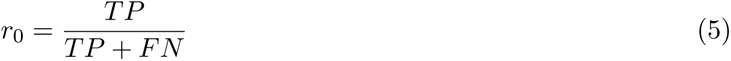

The opposite of recall, is the specificity (*s*_0_), which quantifies the fraction of all negative instances that the classifier identify as positive. Hence, it is known also as the false positive rate (FPR):

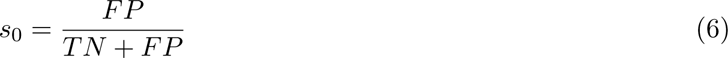

Moreover, the measure of precision is targeted to minimize false positives, and it is also known as the positive predictive value (PPV). It quantifies the fraction of positive predictions that are real:

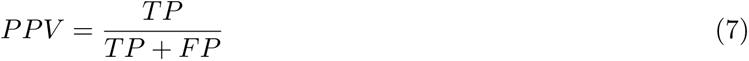

An equivalent measure to precision, but targeted to quantify the fraction of negative predictions that are real, is known as the negative predictive value (NPV):

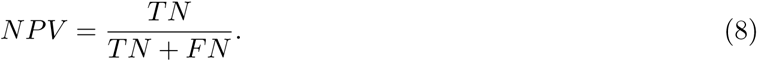

Often, there is a tradeoff between precision and recall. Hence, a quantify known as *F* 1 that effectively combines these two measurements as the harmonic mean of them is defined as:

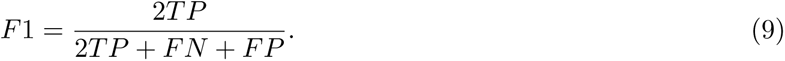

**Table S1:**
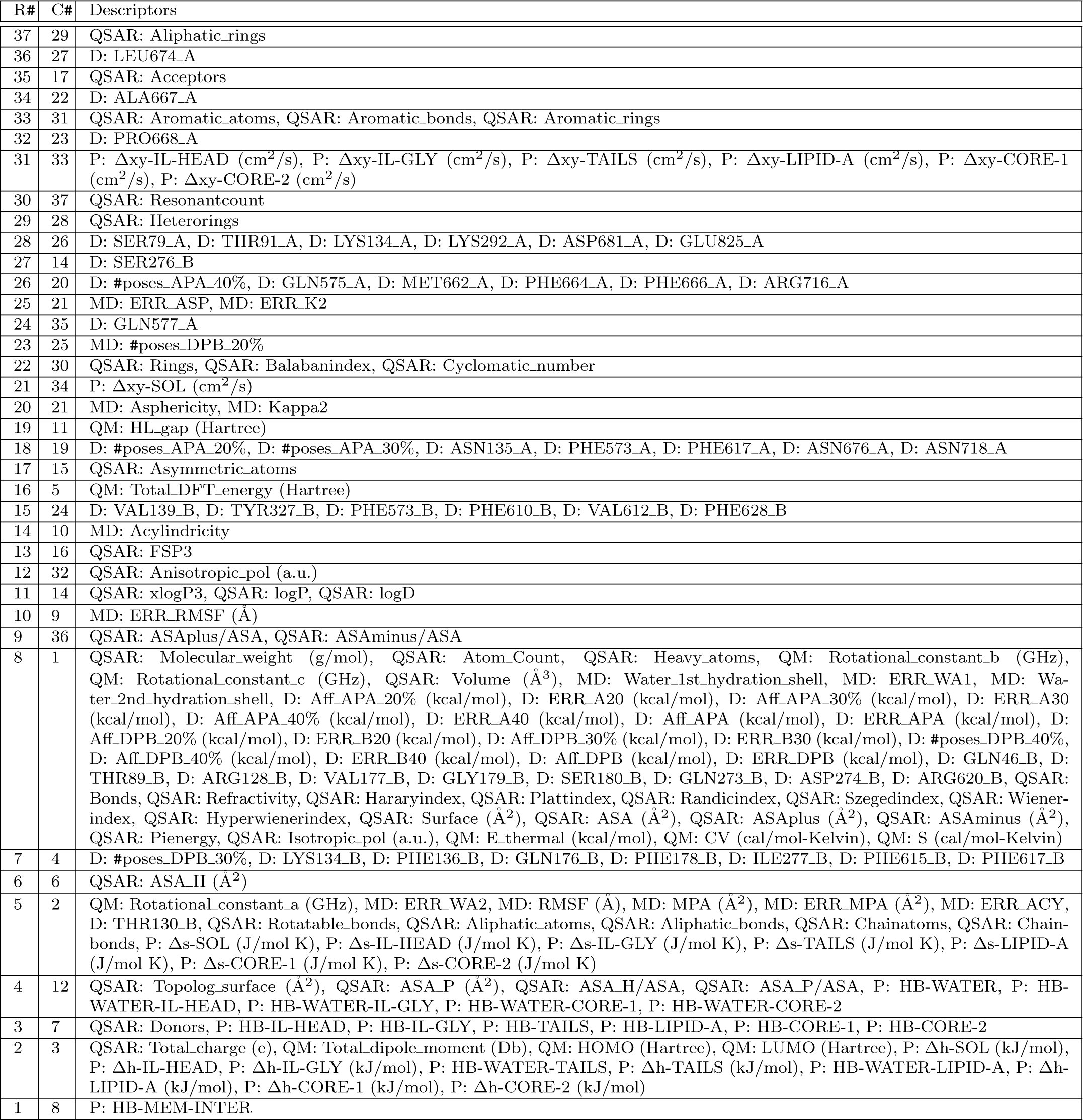
Ranking number (R#) of individual clusters according to permeation predictability, their cluster number (C#), and the descriptors that belong to each of cluster.

**Table S2:** Ranking of clusters *c_j_*up to top-21, where we compare the ordering when using a fixed testing sample, with a ranking produced when the testing sample changes randomly on every single iteration. The ordering of the clusters is robust to these changes for the most part.

| rank | 1 | 2 | 3 | 4 | 5 | 6 | 7 | 8 | 9 | 10 | 11 | 12 | 13 | 14 | 15 | 16 | 17 | 18 | 19 | 20 | 21 |
| --- | --- | --- | --- | --- | --- | --- | --- | --- | --- | --- | --- | --- | --- | --- | --- | --- | --- | --- | --- | --- | --- |
| $c_j$ fixed | 8 | 3 | 4 | 12 | 2 | 6 | 7 | 1 | 36 | 9 | 14 | 32 | 16 | 10 | 24 | 5 | 15 | 19 | 11 | 21 | 34 |
| $c_j$ random | 8 | 3 | 7 | 2 | 12 | 6 | 4 | 1 | 36 | 9 | 13 | 32 | 16 | 10 | 24 | 15 | 19 | 5 | 25 | 11 | 21 |

**Figure S6:**
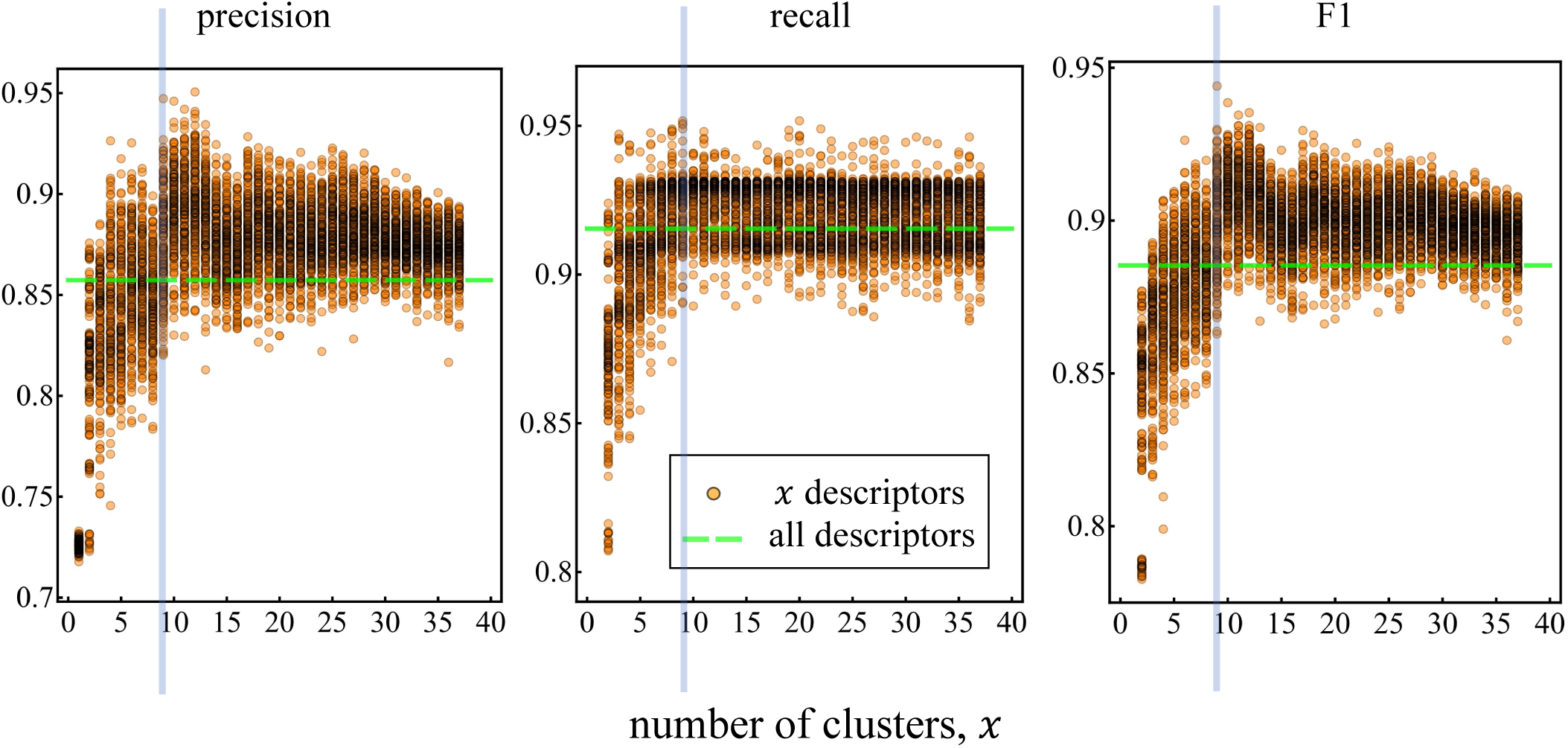
Precision (left), recall (center), and F1 (right) testing results evaluated in the testing portion of the reduction algorithm presented in the main paper. Each orange circle points to the performance of a random combination of *x* compounds, while the horizontal green line is the evaluation metric when all descriptors are considered. Vertical blue line indicates the *x* = 9 mark.

#### 5.2 Evaluation metrics: application

We use these definitions into the classification outcome from our model described in Figs. 4 of the main paper, which also shows the result of accuracy. The results for precision, recall, and F1 for each of the tested combination of *x* descriptors against the testing set of 121 compounds is presented in Fig S6, showing that the cutoff of nine clusters is reasonable in these metrics, as well as, in accuracy.

Figure S7 details the prediction results from the combination listed in Fig. 4(c) in the main paper. The prediction output, which classifies the testing sample of 120 randomly selected molecules as either strong (class 1) or weak (class 0) permeators, is computed over 50 runs of the classification algorithm, where we do random splits of training/validation of the remaining 482 compounds. From the resultant binary output list of 50 outputs, the average target class of each compound is obtained, which could be interpreted as the probability that a given compound to be a strong permeator (class 1). Indeed, if the average output class is near zero, it indicates the algorithm classifies a given compound is a weak permeator (class 0). A direct comparison between our model’s output and the experimental data in the from the IC_50_ ratios is shown in Fig. S7(a), where we see several points of coincidence, as well as, a few misses. Specifically, we see that across the 117 correct classifications of testing compounds as either class 1 or class 0, there are two compounds incorrectly classified as class 1 (FP), and two incorrectly classified as class 0 (FN). Additional model evaluation metrics are presented in Figs. S7(b-c), including the Receiver Operating Characteristic (ROC) curve (solid curve) that is well separated from the limit of random classification (dashed curve). In addition, the evaluation metrics indicates a similar score for recall and for precision, meaning that the algorithm provides a balanced outcome that minimizes almost equally false positives and false negatives. Similarly, the precision or positive predictive value (PPV), and the negative predictive value (NPV), indicates that around 95% of the compounds classified as strong permeators are real, while almost 97% of the compounds classified as weak permeators, are correctly classified as such.

**Figure S7:**
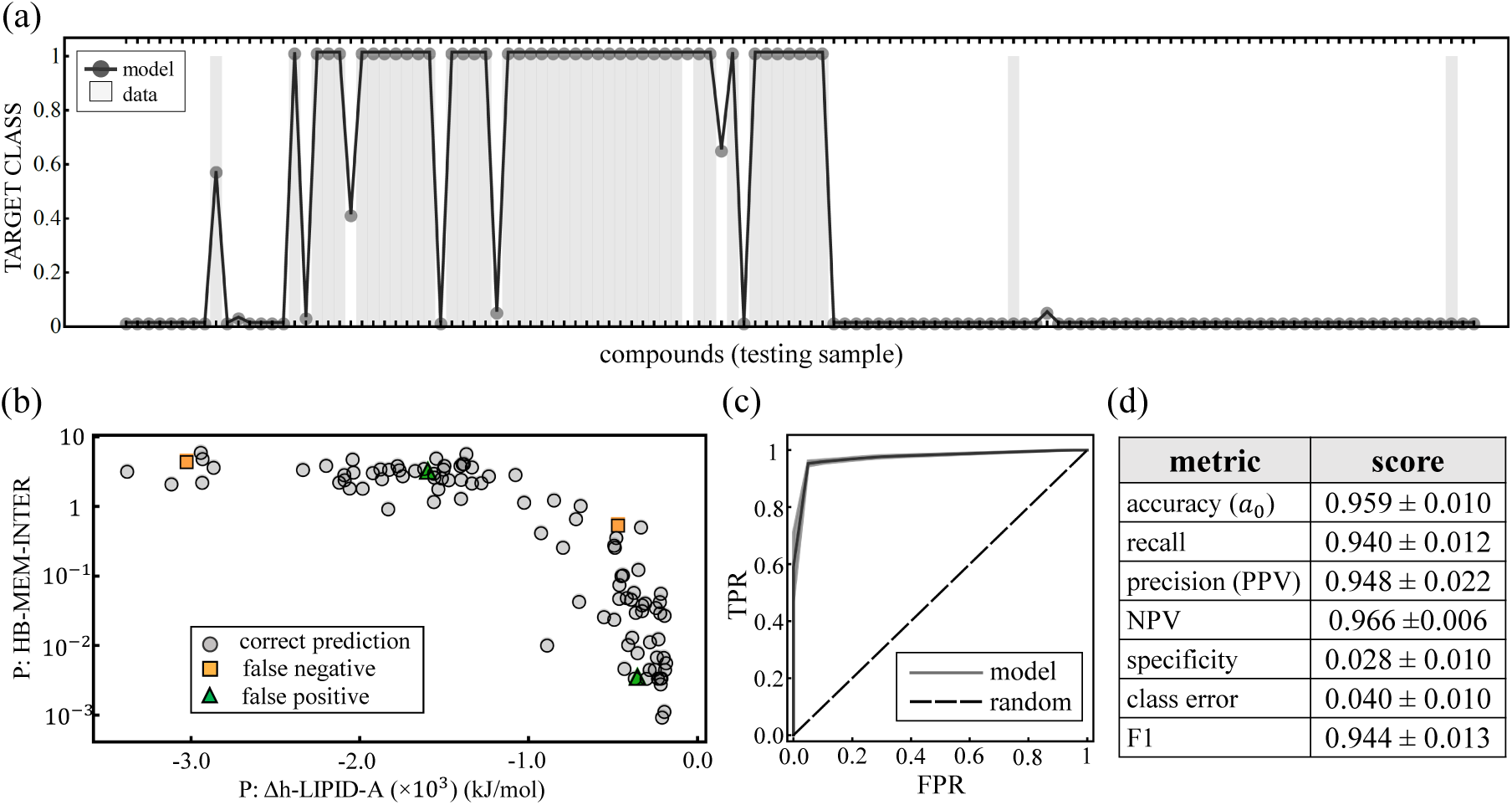
Detailed output of the model in the space of the 120 testing compounds. (a) Direct comparison between the model target class output and the data. The classification output is computed over several random splits of training/validation, hence the average class is reported and could be interpreted as the probability that a given compound is a strong permeator (class 1). (b) Equivalent target class out put but projected in the space of the top-2 descriptors (HB-MEM-INTER, and Δh-LIPID-A). Correct predictions (TP and TN) are shown combined in gray, while the false negatives are orange and the false positive in green. (c) Receiver operating characteristic (ROC) curve associated to our classification model (dark gray) compared to a fully random classification model (black dashed line). (d) Evaluation metrics of the classification model over the testing sample.

#### 5.3 Performance of alternative combination of descriptors from the top-9 clusters

Next, we wonder how far, in terms of performance, other combinations in the top-9 clusters identified by our model are from the optimal. From Fig. 4(c) of the main paper, we see that three of these clusters (6, 8, and 36) have one or two descriptors. This drastically reduces the total number of possible combinations and facilitates a deeper examination of alternative sets of predictors that yield comparable accuracy scores as those presented in Fig. 5 of the main paper. Figure S8 uses the ranking of clusters given by our model (Fig. 4(c) of the main paper) to inspect the top-3, top-5, and top-9 clusters. For the top-3 [Fig.S8(a)] we look at the model performance of cluster 8 with all combinations of pairs from cluster 3 and 7. The top-5 clusters [Fig.S8(b)] uses the optimal descriptors in cluster 8, 7 and 3, and combine them with all the pairs of clusters 2 and 12. Finally, for the top-9 [S8(c)], we use the optimal values of clusters, 2, 3, 7, 12, 6 and 36, with all the pairs from clusters 1 and 4.

Inspecting the top-3 clusters [Fig.S8(a)] we find that permeation descriptors of each cluster tend to perform higher than QM and QSAR descriptors. More specifically, it is found that the enthalpies of the compounds in different regions of the OM (cluster 3), with the exception of CORE 1, pair well with their hydrogen bond interactions (cluster 7). The top-5 shows that the number of hydrogen bonds in the neighboring of the membrane (HB-WATER) performs well when paired with permeation descriptors associated with entropy beyond the one found by the model (Δs-SOL), which is reasonable given the high correlations found within this quantity when measured across the OM [Fig. S4]. Interestingly, it is also found that the polar surface area (ASA P) and some combinations of the polar and hydrophobic surface areas to the water accessible surface area (ASA P/ASA and ASA H/ASA, respectively) also give a good accuracy score and hence could represent alternative descriptors when the entropies across the OM and the hydrogen bonds in water are not known. Finally, the top-9 shows that, in addition to the docking descriptor found by our model (#poses DPB 30%), descriptors quantifying the number of contacts between the compound and residues PHE136 and PHE617 in the DP of MexB give a good prediction performance when compared to descriptors from cluster 1 associated with the size of the compound beyond the Randic index. Hence, these results, in addition to serve as rigorous test of the outcome of our model, deliver alternative combinations of predictors with comparable performance scores to that of the optimal set found by the model that be helpful when screening candidate drugs.

**Figure S8:**
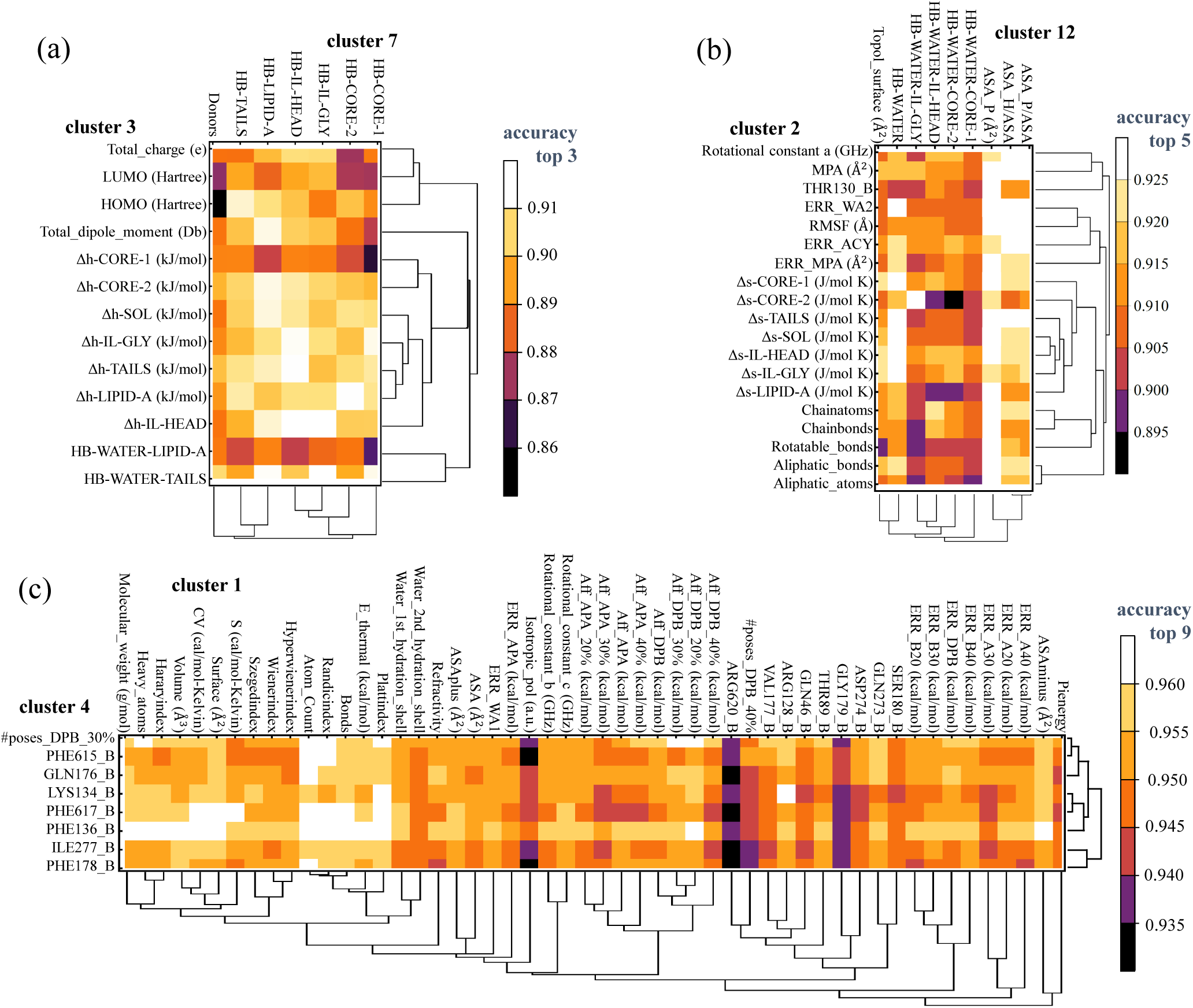
Model performance across clusters. (a) Model’s prediction accuracy using all combinations of descriptors from the top-3 clusters (clusters, 8, 3 and 7). (b) Prediction accuracy for all pairs of descriptors from the forth and fifth ranked clusters together with top-3 descriptors from the optimal combination. (c) Prediction accuracy for all pairs of the seventh and eighth ranked clusters alongside with the top-6 descriptors from the optimal set and descriptor ASAplus/ASA (ranked nineth among clusters).

**Figure S9:**
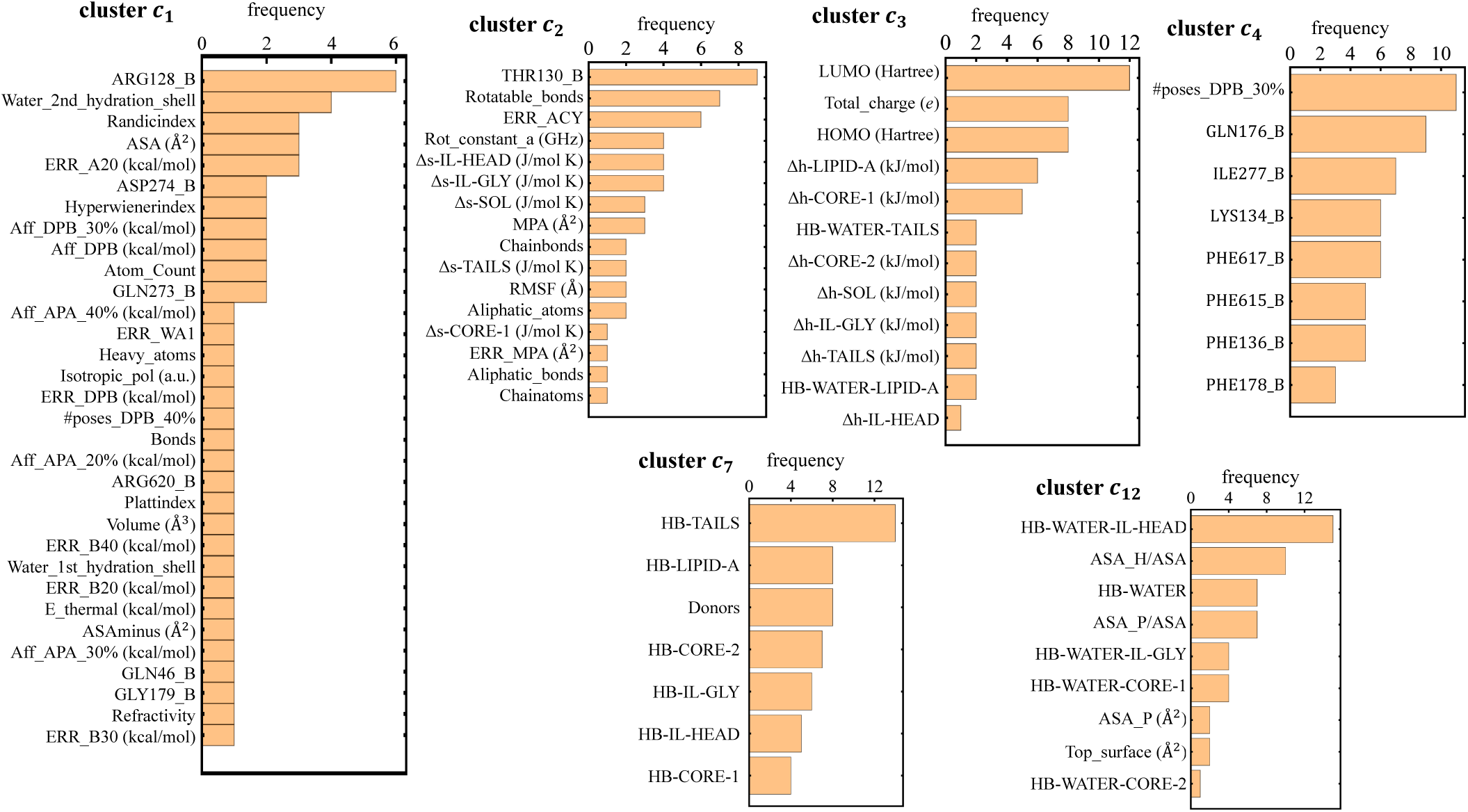
Frequency of identification of specific descriptors for the top-9 clusters across 51 random testing sets of 121 compounds.

### 6 High performance permeation prediction from different combinations of descriptors across testing samples

This section explores additional testing samples using the sampling algorithm described in the main paper. For each random testing set of 120 compounds, 200 different combinations of nine descriptors are randomly chosen (one per cluster), trained and validated in the remaining 482 compounds, and the combination of *x* descriptors that produces the maximum evaluation accuracy in the testing sample is identified. Table S2 lists these descriptors for 50 randomly generated testing samples. A large diversity of combinations is found meaning that the specifics of the data plays a strong role in determining good combinations of descriptors that maximize accuracy.

We further analyze these found combinations by counting the number of times specific descriptors are detected across the different testing samples. The results are shown in the charts of Fig. S9. We find that for the top-9 clusters, there are preferences in choosing some descriptors over other, which highlights their enhanced ability to predict permeation. However, the fact that descriptors are chosen at least once, indicates that for a particular testing sample they play a critical role and therefore its importance cannot be neglected.

To test this finding even further, we compare the accuracy score in the different testing samples for three types of combinations of descriptors. We call combination *j*, to the set of descriptors that yield the highest accuracy for the testing set *j*. Thus, if we are working with 50 testing samples, each of them would have a particular combination of descriptors given by the Table S2. By contrast, we call combination *a* the set of descriptors that is found for one single testing sample and use it on the different testing samples. For the case shown in Fig. S10, we use the last row of Table S2, which is also the combination listed in Fig. 4(c) of the main paper. Finally, we use the most frequent descriptor for each cluster as indicated in Fig. S9. This is referred to as combination *b*. Figure S10 illustrates the results for these three contrasting cases, where we find that the curves tend to follow each other very well and hence the differences between these scores are rather small. Indeed, in most instances, combination *a* tends to be slightly below the other two cases, but the calculating the average difference with combination *j* we find that it is only 2.33%, while combination *b* has an average difference of 1.3% with combination *j*. Hence, the information provided by the descriptors of the different combinations is highly correlated with the IC_50_ ratios, and it leaves a very small room for improvement or optimization.

**Table S3:**
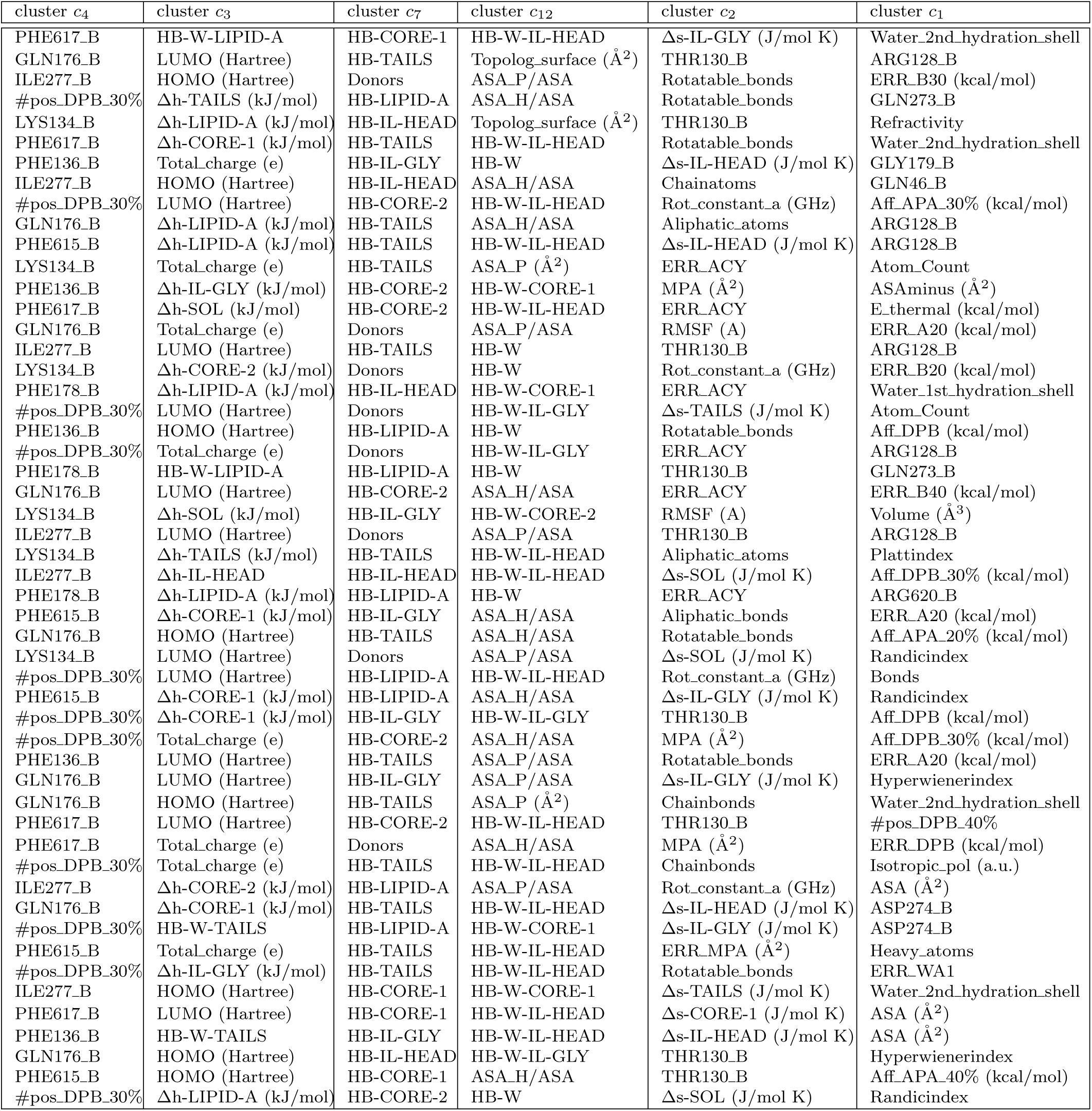
Example of the variety of combinations of descriptors from the top-9 clusters identified that lead to maximal accuracy for different random testing samples. Since three clusters in the top-9 contain only one (*c*_8_ and *c*_6_) or two descriptors (*c*_36_), the table lists those for clusters of size greater than 2 only.

**Figure S10:**
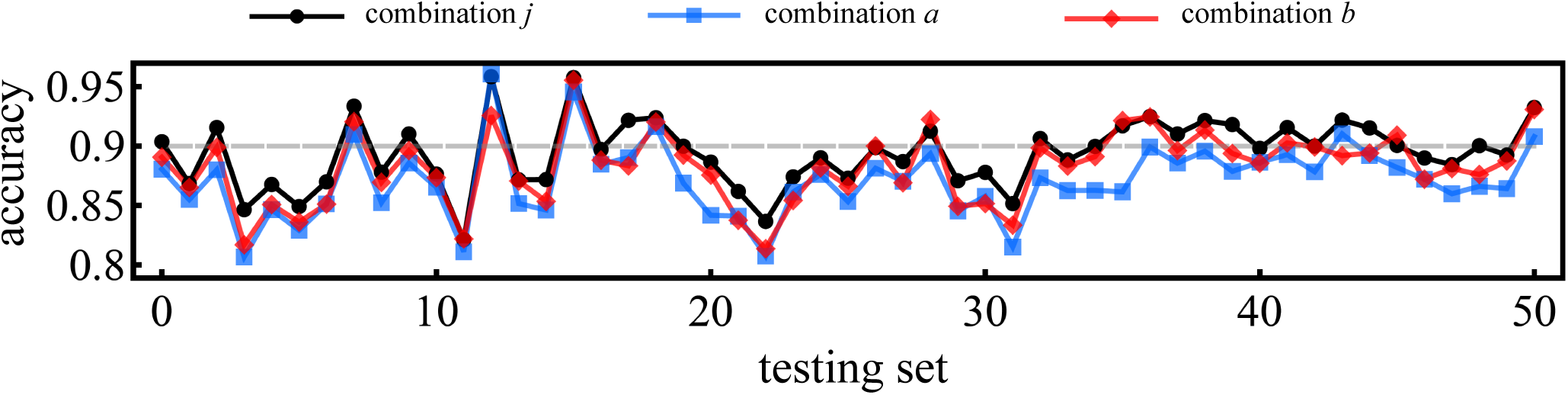
Accuracy score over 50 randomly assembled testing samples using the different combinations of descriptors as explained in the text: combination *j*, combination *a*, and combination *b*. Gray horizontal line indicates the 90% mark in accuracy.

### 7 Classification of active compounds according to predictability

The analysis performed in section 2.4 is further examined in order to extract insights about the predictability of the different compounds.

#### 7.1 Permeation class threshold based on empirical IC_50_ ratio

The IC_50_ ratio threshold separating the permeation classification of the compounds presented in the main manuscript is 0.5. This choice comes from extensive calculations with several choices of thresholds. For each choice we have run our reduction algorithm over the entire set of clusters of descriptors where we identified the combination that maximizes the classification performance (Fig 4(a) in the main manuscript). Subsequently, we test these descriptors across 100 randomly constructed testing samples of 120 compounds each where we identify the groups G (always predicted correctly), R (always predicted incorrectly), and B (sometimes predicted correctly and sometimes incorrectly) for each ratio threshold tested (Fig. 6(a) in the main manuscript). Figure S11 shows that a ratio of 0.5 maximizes the number of compounds predicted correctly (set G on top) at every run, while simultaneously minimizing the number of compounds predicted incorrectly (set R in the middle) and those producing mixed results (set B in the bottom). As shown, other threshold choices produces higher classification errors and larger fluctuations.

**Figure S11:**
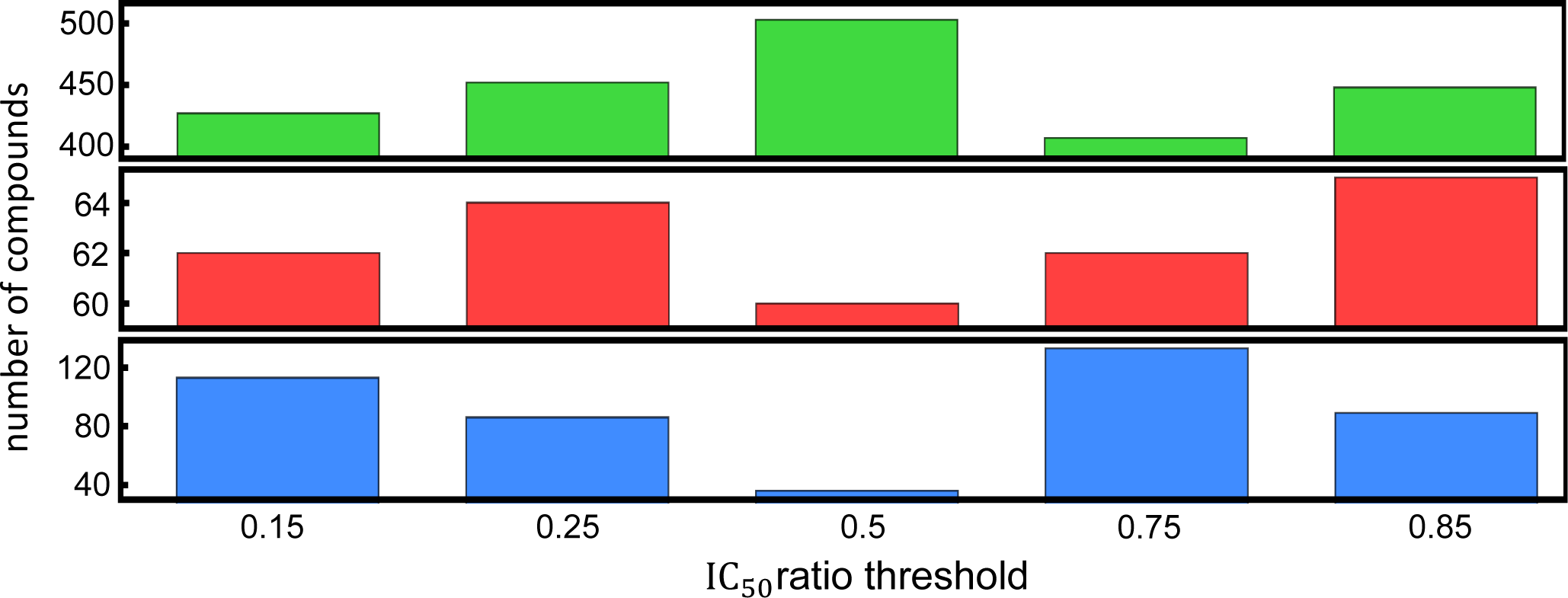
Active compounds classification according to predictability for different choices of IC_50_ ratio threshold. The size of the sets G, R and B, are shown in the top, middle, and bottom panel, respectively. A threshold of 0.5 maximizes the compounds in the set G while minimizing the sets R and B, as shown.

**Table S4:** Comparison of the different evaluation metrics for different combination of predictability sets: all sets, G+R, G+B, and R+B. For each combination set we show the average metric across 100 randomly assembled testing samples.

| metric/set | all | G + B | G + R | R + B |
| --- | --- | --- | --- | --- |
| accuracy | 0.868±0.027 | 0.969±0.016 | 0.893±0.027 | 0.720±0.086 |
| recall | 0.820±0.051 | 0.971±0.027 | 0.843±0.044 | 0.721±0.132 |
| precision | 0.853±0.047 | 0.951±0.032 | 0.893±0.042 | 0.756±0.130 |
| NPV | 0.878±0.037 | 0.981 ±0.017 | 0.893±0.031 | 0.698±0.143 |
| specificity | 0.097±0.034 | 0.031±0.021 | 0.070±0.029 | 0.266±0.142 |
| class error | 0.131±0.027 | 0.030±0.016 | 0.106±0.027 | 0.279±0.086 |
| F1 | 0.835±0.034 | 0.960±0.022 | 0.867±0.035 | 0.726±0.092 |

#### 7.2 Parameter regions associated with the different permeation classes

Focusing on the threshold of 0.5 and for the set of nine descriptors highlighted by the reduction algorithm (see Fig 4 in the main paper), we examine the two-dimensional scatter plots of the compounds of the set G. This yield a total of 36 scatter plots that are shown in Fig. S12. Interestingly, there are clear regions that separate strong (red) and weak (blue) permeators. The set G contain the largest amount of compounds (503), accounting for 83% of all active molecules analyzed. This finding allow us to draw simple rules of permeation based on the descriptor values associated with a given compound. For comparison, Table S4 provides the evaluation metric for different combinations of predictability sets: all, G+B, G+R, and R+B.

In order to compare this behavior of the set G with the compounds of the sets R and B, Fig. S13 illustrates these three sets in the space of the top-2 descriptors that better predict their permeation properties. Similar to the result of the set G, compounds in the set R are also separated into specific regions in the parameter space. However, the regions associated to weak and strong permeators are opposite to those in the compounds of the set G. This explains why they are always missed by the prediction algorithm. Finally, the set B has compounds some of which belong to the pattern shown by the set G and the remaining show the pattern presented in the set R.

**Figure S12:**
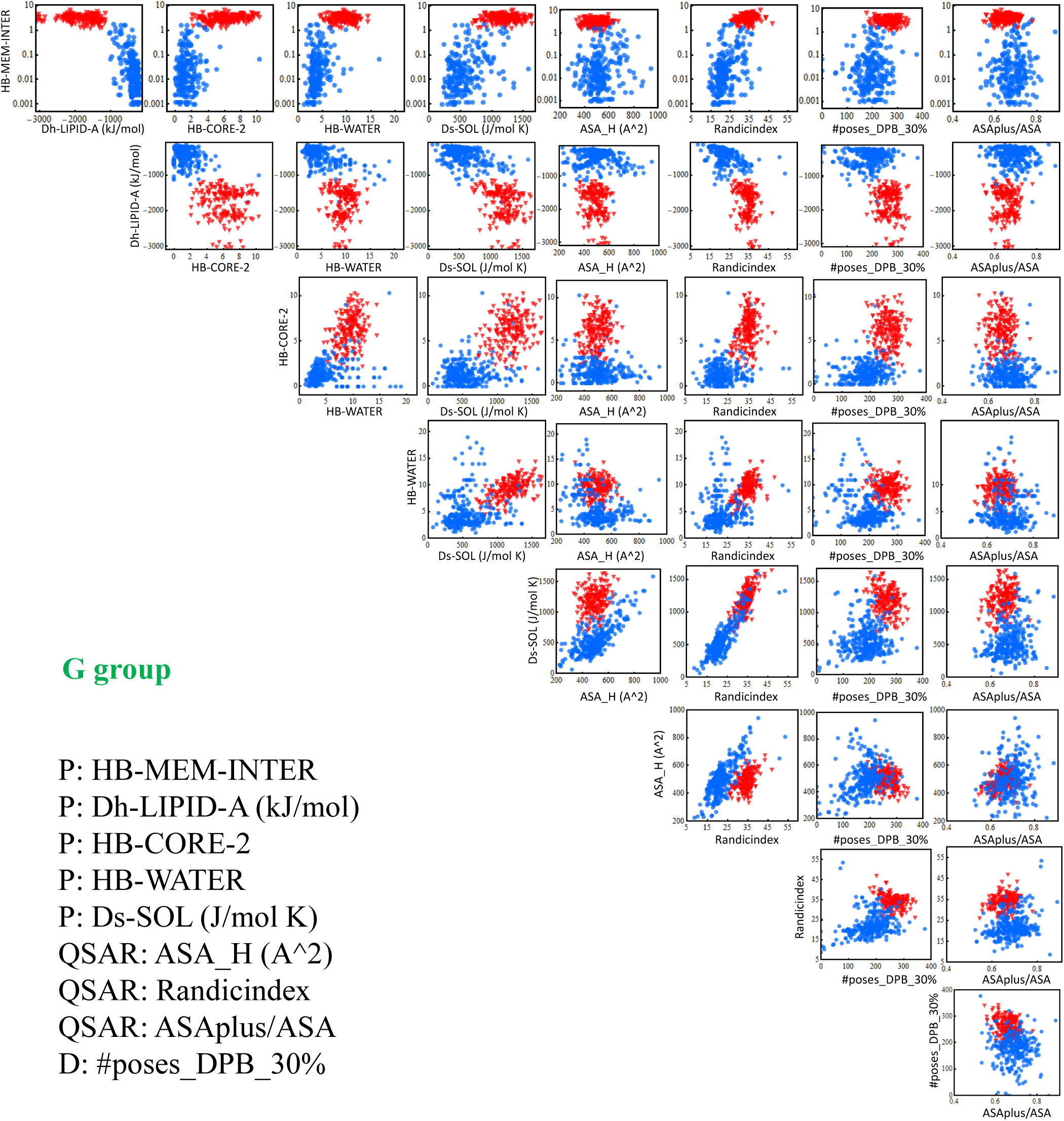
Compounds of the set G projected over the nine descriptors highlighted by the reduction algorithm, as listed in the bottom left. In each of the scatter plots shown, red dots represent strong permeators, while blue are weak permeators. For each row (column) the y-(x-)axis represents a single descriptor, as shown.

**Figure S13:**
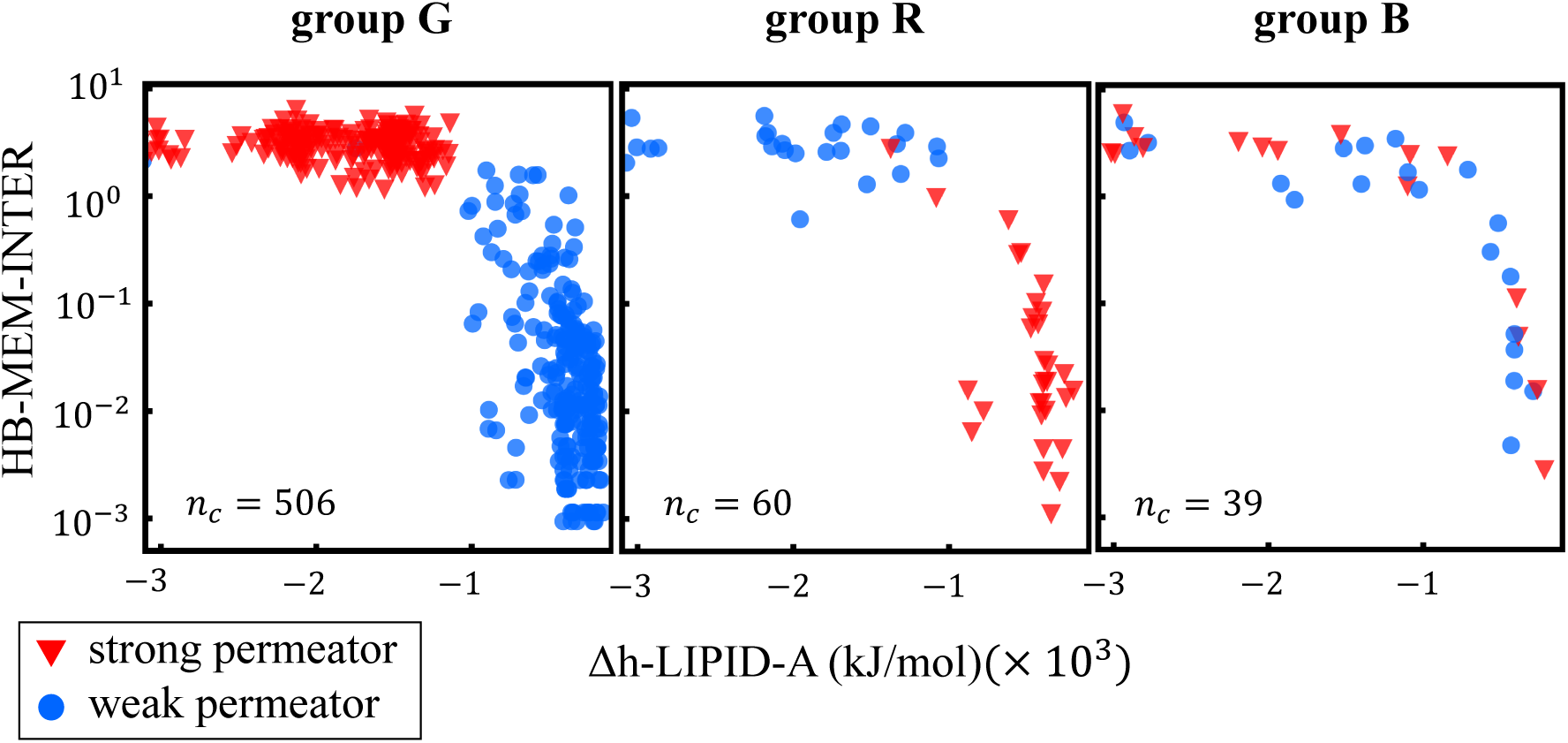
Projection of the 605 active compounds over the top-2 descriptors that predict permeation the best for the three predictive classification groups: set G (left panel), set R (central panel), and set B (right panel).

#### 7.3 Structural aspects of compounds in critical subgroups

A classification of the compounds by means of a complete Tanimoto similarity analysis reveals 233 subgroups. As explained in the main manuscript, around 90% of the compounds in sets R and B are concentrated in 10 Tanimoto subgroups. Figure S14 shows the results of our modeling classification in the seven subgroups not shown in Figure 5(d) of the main manuscript. Our analysis identifies descriptors able to separate weak and strong permeators for some of these subgroups including some compounds of the sets R and B. For other subgroups, however, it remains a challenge. The structural features that distinguish these 10 subgroups are detailed below:

- SB71 are the NSC-33353 analogs made as EPI’s. Most of those compounds had a substituted naphthyl amide group present. This is the small subset that didn’t have a naphthyl amide but instead had phenyl or biphenyl. So, this seems clear relative to the other compounds in the NSC-33353 series. What is also interesting, it is these phenyl and biphenyl are currently being made because they work in WT Acinetobacter. But not E. coli or PAO1.
- SB117 is full of NSC-60339 analogs. Unlike the subgroup below, these typically have 5-aromatic rings in them or more and tend to be more symmetrical with 2 or more of the dihydroimidazoline groups. So, this subgroup is mostly the original hits and analogs. The subgroup below (SB118) is typically 2-3-aromatic and non-symmetrical with only one dihydroimiazoline group. So, that’s the biggest difference. This subgroup is the analogs with more than one dihydroimidazoline moiety in them. They are going to have larger MW compared to SB118 as well.
- SB118 is composed of the NSC-60339 analogs we got from the NCI and synthesized to generate new EPI’s. They all have that same group (4-(4,5-dihydro-1H-imidazol-2-yl)aniline) that all of the NSC-60339 analogs have in it. These are generally fairly linear with only a few rotatable bonds and generally fairly lipophilic.
- SB112 is a mix of the original NSC-125028 compounds (MUKB inhibitors) and NSC-60339 (Original EPI compounds). There is a mix of charged and uncharged compounds. Finally, analogs and the intermediates to make them. They are all polyaromatic with amide linkers. There are not many rotatable bonds, generally lipophilic.
- SB167 is an amide derived from 3-aminoquinoline with one exception is cmpd, which is not a quinoline but a naphthyl. It could have been in this group or in SB168. The reason it was not could be that most, but not all, of SB167 also has a formal +1 charge while most of SB168 does not.
- SB168 all have an amide derived from a 6-aminoquinoline group present in their structure. There were 3 exceptions. One of them has a 6-substituted quinoline in it, but it is not an amide. And then two compounds which do not have the quinoline but instead have a cyclic boronate derived amide in them.
- SB169 has the phenyl ethyl side chain (from the unnatural amino acid) as opposed to the phenyl alanine or substituted phenyl alanine, that is present in most of the others.
- 12/13 compounds of SB170 contain a phenyl boronic acid group. In general, these are less rigid and lower MW than the previous Rempex subgroups discussed. The one non-boronic acid has a phenol instead but otherwise does look somewhat similar and the PKA’s of phenol and boronic acid are somewhat close. This subgroup is somewhat related structurally to SB223. More so than the other Rempex subgroups.
- SB201 are benzothiazoles (BTZs) from JKW lab. The majority of the BTZ’s made had a n-propylimidazole group on them. That was key for the antibacterial activity but also made them substrates for efflux and contributed to poor PK. A handful of them were made without the n-propylimidazole group to try and change the properties and hopefully improve their permeability etc. That small subset is captured in subgroup SB201. the n-propylimidazole group was replaced by simple heterocyclic groups such as piperidine, pyrrolidine etc.
- SB223 are Rempex compounds. With one exception none of these compounds are charged (N-methylated). There is also one boronic acid in this group. They are also more in general have more SP3 character. MW is probably lower on average compared to the other Rempex subgroups, but certainly less rigid, not as elaborated structurally and not formally charged.

**Figure S14:**
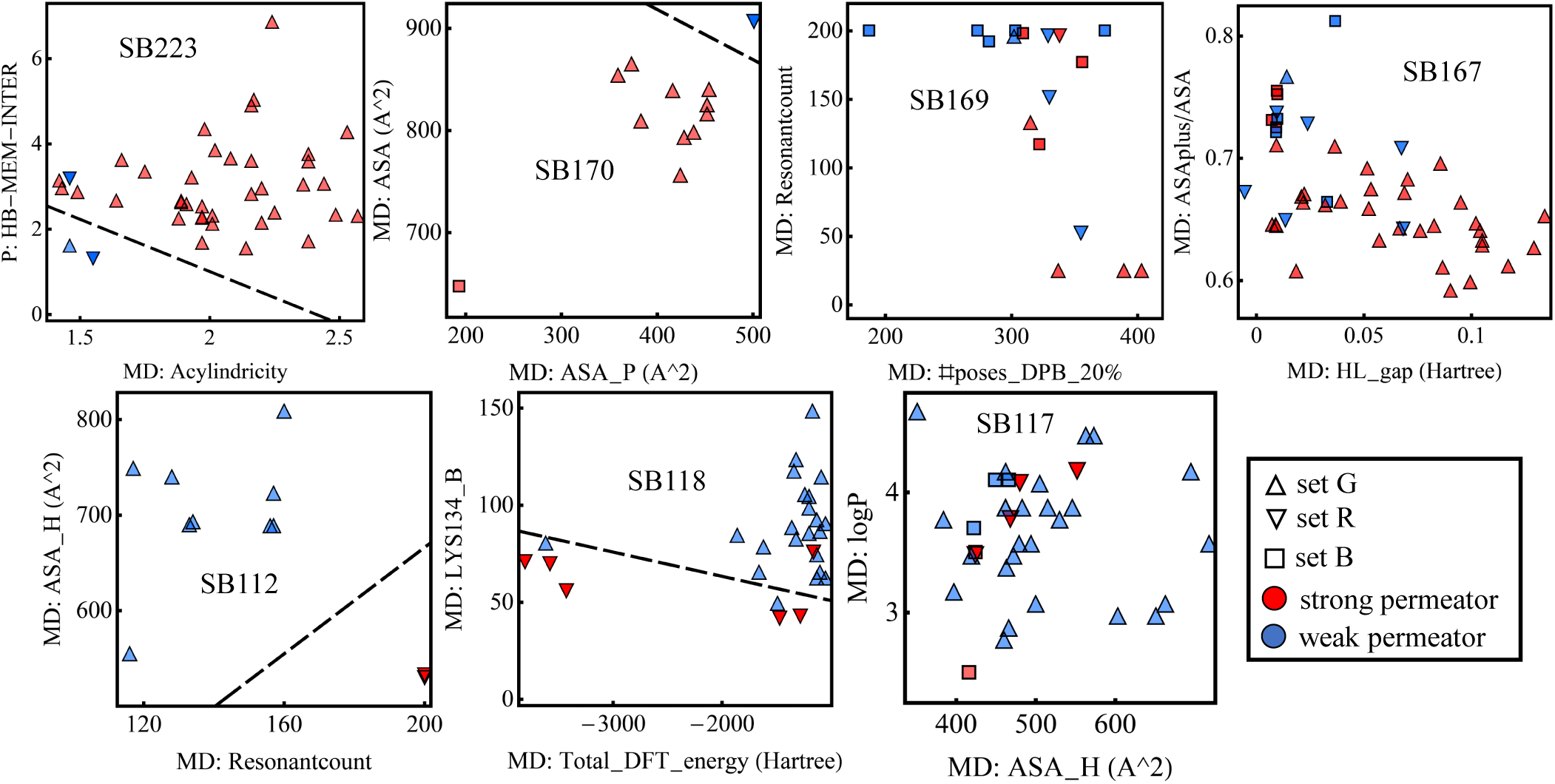
Complementary analysis of relevant descriptors for the remaining seven Tanimoto subgroups that contain relevant number of compound of sets R and B, as illustrated in Figure 5(d) of the main manuscript. In cases where the separation between weak and strong permeators is somewhat clear, it is marked by a class boundary black line found using an SVM algorithm.

### 8 Descriptor ranges associated to weak and strong permeation

The consistency of the patterns found for the largest group of compounds (set G), allows us to draw simple rules of permeation based on a compound’s parameter values. To this end, we use the top-9 clusters that better correlate with the IC_50_ ratios. This results in a total of 112 descriptors. Tables S5 and S6 shows the ranking of these descriptors that better correlate with permeation individually. In these tables we compare the metrics for the set G with that of all of the active compounds. The ranking is according to accuracy in the latter set. The value *t_c_*is the critical threshold dividing strong and weak permeators according to column L|R. For example, when looking at descriptor *δ_j_*, if *δ_j_ > t_c_*, its classification would be that of R (i.e., W or S for weak and strong permeator, as listed in the L|R column), and it is L, otherwise. The threshold is estimated using a Support Vector Machine [38] (SVM) algorithm with linear kernel that separates the strong and weak permeators data using a maximum-margin hyperplane. For the one-dimensional case (i.e., one descriptor), *t_c_* is a single point. Extensions to two and three dimensions are also implemented for comparison, where the SVM find the optimal line [Fig. S12] or plane [Fig. S13] to separate the target classes, respectively.

A similar analysis now using sets of two- and three-descriptors is also performed, where we trained a 2- and 3-dimensional support vector machine on all combinations of two and three descriptors of the top-9 clusters (no more than one descriptor per cluster per combination), respectively. Accordingly, we found an enhancement in the performance accuracy for different combinations of two and three descriptors. For simplicity, we have use a straight line and a plane for the cut off of the two- and three-descriptor analysis, respectively. The top-40 combinations and corresponding parameters are listed in the Tables S7 and S8, while we highlight four sets of three-descriptors combinations that classifies well strong and weak permeators (see Fig. S16).

The two-descriptor analysis highlights the role of the electrostatic properties and the electronic structure, together with, the hydrophobic and polar area, of the compounds [Fig. S15]. These pairs of descriptors yield accuracy scores up to 88.1% computed across the entire set of active compounds. In addition, we find several pairs of descriptors comprised by one of permeation (either HB or enthalpy) and one of docking (#poses or contacts to key residues in DP of MexB), or permeation with QM or QSAR descriptors. Finally, the three-descriptor analysis finds the QSAR descriptor quantifying the ratio of the hydrophobic area to the water accessible surface area (i.e., ASA H/ASA) in myriad combinations with either one permeation and one docking descriptor, or one docking with another QSAR or QM. This separation talks about an interesting relationship among the different types of descriptors and how their information can be harnessed in order to better identify strong and weak permeators. Some examples are found in Fig. S16 reaching accuracy scores of 88.42% obtained across the whole set of active compounds.

**Table S5:**
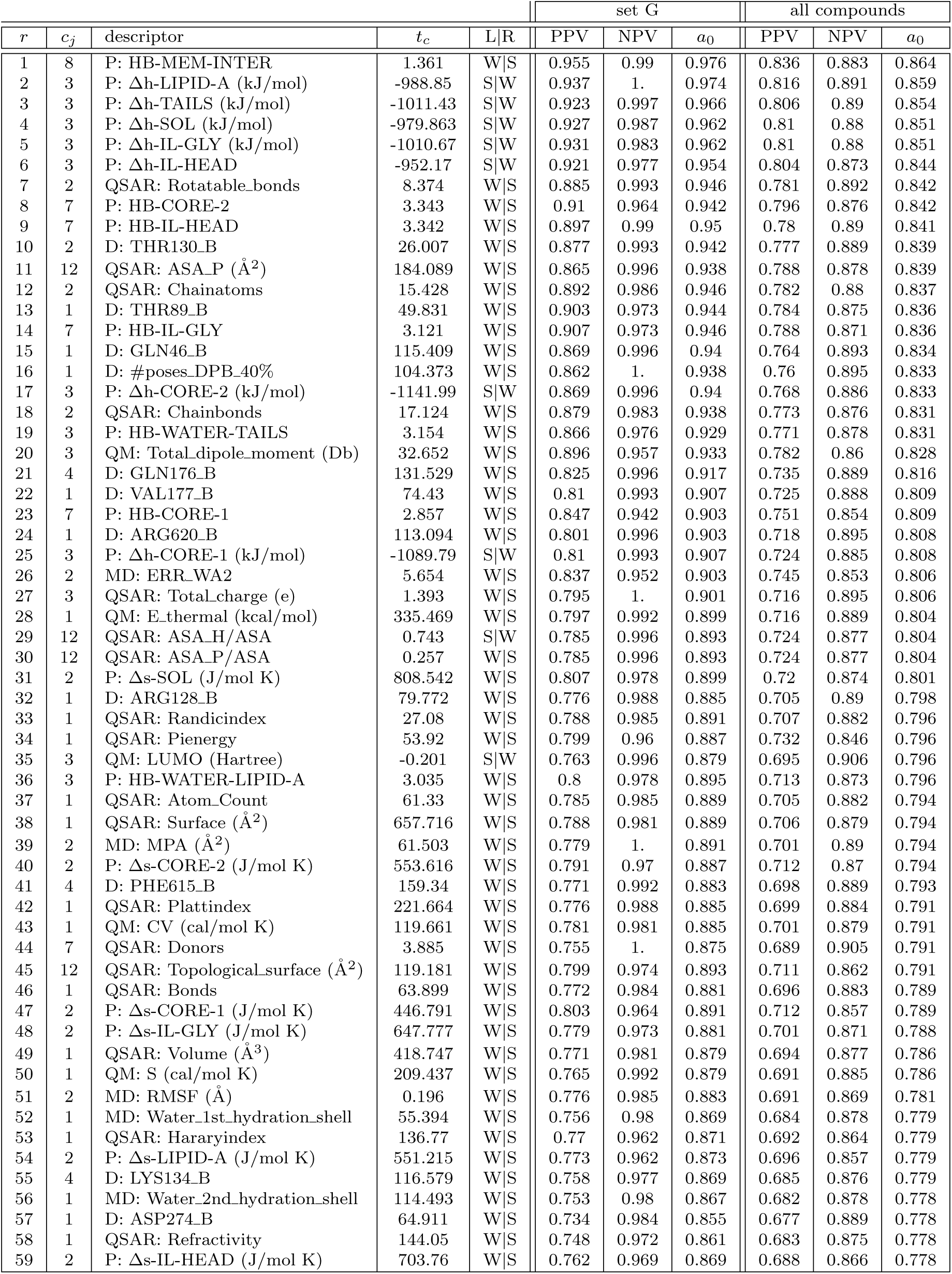
Individual descriptor ranges associated with strong (SP) and weak (WP) OM permeation for the top-60 descriptors according to the accuracy score of all active compounds. For each ranking descriptor *r* belonging to cluster *c_j_*, the threshold *t_c_* separating the target classes, is listed in the measurement units of the descriptor. Column L|R indicates whether WP are associated with descriptor values smaller than *t_c_* and hence SP associated with values larger than *t_c_* (entry W|S), or vice versa (entry S|W). The evaluation metrics of positive predictive value (PPV), negative predictive value (NPV) and accuracy (*a*_0_) are listed for each descriptor for the compounds of the set G only and for all active compounds, as indicated.

**Table S6:** Individual descriptor ranges associated with strong (SP) and weak (WP) OM permeation for the descriptors below the top-60 according to the accuracy score of all active compounds. For each ranking descriptor *r* belonging to cluster *c_j_*, the threshold *t_c_* separating the target classes, is listed in the measurement units of the descriptor. Column L|R indicates whether WP are associated with descriptor values smaller than *t_c_* and hence SP associated with values larger than *t_c_* (entry W|S), or vice versa (entry S|W). The evaluation metrics of positive predictive value (PPV), negative predictive value (NPV) and accuracy (*a*_0_) are listed for each descriptor for the compounds of the set G only and for all active compounds, as indicated.

| $r$ | $c_j$ | descriptor | $t_c$ | L\|R | set G | | | all compounds | | |
| --- | --- | --- | --- | --- | --- | --- | --- | --- | --- | --- |
| | | | | | PPV | NPV | $a_0$ | PPV | NPV | $a_0$ |
| 60 | 1 | QSAR: Heavy_atoms | 33.457 | W\|S | 0.764 | 0.966 | 0.869 | 0.687 | 0.863 | 0.776 |
| 61 | 1 | MD: ERR_WA1 | 4.355 | W\|S | 0.771 | 0.941 | 0.863 | 0.696 | 0.847 | 0.776 |
| 62 | 1 | QSAR: Wienerindex | 4272.3 | W\|S | 0.777 | 0.942 | 0.867 | 0.696 | 0.847 | 0.776 |
| 63 | 1 | D: SER180_B | 52.181 | W\|S | 0.749 | 0.965 | 0.859 | 0.682 | 0.867 | 0.774 |
| 64 | 1 | QSAR: Hyperwienerindex | 23670. | W\|S | 0.788 | 0.92 | 0.863 | 0.704 | 0.832 | 0.774 |
| 65 | 1 | QSAR: Molecular_weight (g/mol) | 473.818 | W\|S | 0.755 | 0.965 | 0.863 | 0.681 | 0.864 | 0.773 |
| 66 | 2 | MD: ERR_MPA ($\text{\AA}^2$) | 5.67 | W\|S | 0.749 | 0.976 | 0.863 | 0.679 | 0.869 | 0.773 |
| 67 | 2 | MD: ERR_ACY | 0.396 | W\|S | 0.758 | 0.977 | 0.869 | 0.68 | 0.867 | 0.773 |
| 68 | 2 | QSAR: Aliphatic_atoms | 19.48 | W\|S | 0.748 | 0.984 | 0.865 | 0.676 | 0.874 | 0.773 |
| 69 | 12 | P: HB-WATER-IL-GLY | 3.726 | W\|S | 0.761 | 0.941 | 0.857 | 0.693 | 0.844 | 0.773 |
| 70 | 1 | D: GLN273_B | 39.855 | W\|S | 0.743 | 0.976 | 0.859 | 0.678 | 0.866 | 0.771 |
| 71 | 1 | D: ERR_A30 (kcal/mol) | 0.603 | W\|S | 0.805 | 0.891 | 0.857 | 0.719 | 0.801 | 0.768 |
| 72 | 2 | P: Δs-TAILS (J/mol K) | 601.65 | W\|S | 0.758 | 0.955 | 0.861 | 0.682 | 0.849 | 0.768 |
| 73 | 12 | P: HB-WATER-CORE-1 | 3.511 | W\|S | 0.771 | 0.931 | 0.859 | 0.695 | 0.828 | 0.768 |
| 74 | 1 | D: ERR_APA (kcal/mol) | 0.6 | W\|S | 0.77 | 0.906 | 0.847 | 0.7 | 0.818 | 0.766 |
| 75 | 12 | P: HB-WATER | 6.742 | W\|S | 0.733 | 0.949 | 0.843 | 0.67 | 0.851 | 0.761 |
| 76 | 1 | QSAR: ASA ($\text{\AA}^2$) | 684.798 | W\|S | 0.742 | 0.939 | 0.845 | 0.67 | 0.841 | 0.758 |
| 77 | 3 | QM: HOMO (Hartree) | -0.305 | S\|W | 0.698 | 0.991 | 0.831 | 0.644 | 0.914 | 0.758 |
| 78 | 1 | D: ERR_A20 (kcal/mol) | 0.627 | W\|S | 0.763 | 0.905 | 0.843 | 0.685 | 0.813 | 0.756 |
| 79 | 7 | P: HB-TAILS | 2.122 | W\|S | 0.767 | 0.888 | 0.837 | 0.691 | 0.805 | 0.756 |
| 80 | 1 | D: ERR_A40 (kcal/mol) | 0.556 | W\|S | 0.721 | 0.938 | 0.831 | 0.664 | 0.843 | 0.755 |
| 81 | 2 | QSAR: Aliphatic_bonds | 21.404 | W\|S | 0.716 | 0.979 | 0.841 | 0.654 | 0.867 | 0.755 |
| 82 | 1 | D: ERR_DPB (kcal/mol) | 0.735 | W\|S | 0.757 | 0.89 | 0.833 | 0.685 | 0.802 | 0.751 |
| 83 | 1 | D: GLY179_B | 73.627 | W\|S | 0.709 | 0.955 | 0.829 | 0.649 | 0.868 | 0.751 |
| 84 | 1 | QSAR: Szegedindex | 5787.5 | W\|S | 0.743 | 0.916 | 0.837 | 0.671 | 0.82 | 0.751 |
| 85 | 7 | P: HB-LIPID-A | 1.825 | W\|S | 0.777 | 0.877 | 0.837 | 0.686 | 0.796 | 0.75 |
| 86 | 1 | QSAR: ASAMinus ($\text{\AA}^2$) | 221.86 | W\|S | 0.726 | 0.928 | 0.831 | 0.662 | 0.823 | 0.746 |
| 87 | 1 | D: ERR_B40 (kcal/mol) | 0.733 | W\|S | 0.724 | 0.895 | 0.817 | 0.658 | 0.806 | 0.738 |
| 88 | 4 | D: #poses_DPB_30% | 214.971 | W\|S | 0.692 | 0.958 | 0.817 | 0.635 | 0.859 | 0.738 |
| 89 | 1 | D: ERR_B20 (kcal/mol) | 0.726 | W\|S | 0.721 | 0.879 | 0.81 | 0.661 | 0.798 | 0.736 |
| 90 | 1 | D: Aff_APA_30% (kcal/mol) | -9.719 | S\|W | 0.697 | 0.958 | 0.821 | 0.635 | 0.846 | 0.735 |
| 91 | 1 | D: Aff_DPB_20% (kcal/mol) | -10.321 | S\|W | 0.68 | 0.978 | 0.813 | 0.629 | 0.866 | 0.735 |
| 92 | 1 | D: Aff_DPB_40% (kcal/mol) | -10.393 | S\|W | 0.674 | 0.982 | 0.81 | 0.625 | 0.871 | 0.733 |
| 93 | 1 | D: Aff_DPB_30% (kcal/mol) | -10.361 | S\|W | 0.674 | 0.982 | 0.81 | 0.624 | 0.868 | 0.731 |
| 94 | 12 | P: HB-WATER-IL-HEAD | 4.076 | W\|S | 0.699 | 0.898 | 0.804 | 0.648 | 0.806 | 0.731 |
| 95 | 1 | D: Aff_APA (kcal/mol) | -9.447 | S\|W | 0.663 | 0.952 | 0.794 | 0.625 | 0.851 | 0.728 |
| 96 | 1 | D: ERR_B30 (kcal/mol) | 0.737 | W\|S | 0.698 | 0.888 | 0.8 | 0.644 | 0.803 | 0.728 |
| 97 | 4 | D: PHE617_B | 95.925 | W\|S | 0.681 | 0.93 | 0.802 | 0.631 | 0.832 | 0.728 |
| 98 | 1 | D: Aff_APA_40% (kcal/mol) | -9.665 | S\|W | 0.665 | 0.977 | 0.802 | 0.619 | 0.866 | 0.726 |
| 99 | 1 | D: Aff_DPB (kcal/mol) | -10.28 | S\|W | 0.667 | 0.982 | 0.804 | 0.618 | 0.869 | 0.726 |
| 100 | 1 | D: Aff_APA_20% (kcal/mol) | -9.634 | S\|W | 0.663 | 0.944 | 0.792 | 0.616 | 0.846 | 0.72 |
| 101 | 1 | QSAR: ASAPlus ($\text{\AA}^2$) | 462.964 | W\|S | 0.678 | 0.892 | 0.788 | 0.619 | 0.799 | 0.71 |
| 102 | 4 | D: PHE136_B | 123.694 | W\|S | 0.632 | 0.944 | 0.766 | 0.591 | 0.837 | 0.697 |
| 103 | 2 | QM: Rotational_constant_a (GHz) | 0.324 | S\|W | 0.614 | 1. | 0.758 | 0.577 | 0.876 | 0.688 |
| 104 | 12 | P: HB-WATER-CORE-2 | 3.906 | W\|S | 0.65 | 0.844 | 0.754 | 0.604 | 0.756 | 0.687 |
| 105 | 1 | QM: Isotropic_pol (a.u.) | 406.648 | W\|S | 0.634 | 0.827 | 0.738 | 0.592 | 0.759 | 0.68 |
| 106 | 4 | D: ILE277_B | 123.369 | W\|S | 0.605 | 0.855 | 0.724 | 0.58 | 0.783 | 0.677 |
| 107 | 1 | QM: Rotational_constant_c (GHz) | 0.053 | S\|W | 0.586 | 0.967 | 0.724 | 0.555 | 0.863 | 0.663 |
| 108 | 4 | D: PHE178_B | 176.99 | W\|S | 0.575 | 0.863 | 0.7 | 0.558 | 0.784 | 0.657 |
| 109 | 1 | QM: Rotational_constant_b (GHz) | 0.066 | S\|W | 0.533 | 0.948 | 0.661 | 0.52 | 0.86 | 0.62 |
| 110 | 36 | QSAR: ASAPlus/ASA | 0.677 | S\|W | 0.5 | 0.742 | 0.615 | 0.502 | 0.699 | 0.594 |
| 111 | 36 | QSAR: ASAMinus/ASA | 0.323 | W\|S | 0.5 | 0.742 | 0.615 | 0.502 | 0.699 | 0.594 |
| 112 | 6 | QSAR: ASA_H ($\text{\AA}^2$) | 500.76 | S\|W | 0.391 | 0.622 | 0.5 | 0.423 | 0.608 | 0.512 |

**Figure S15:**
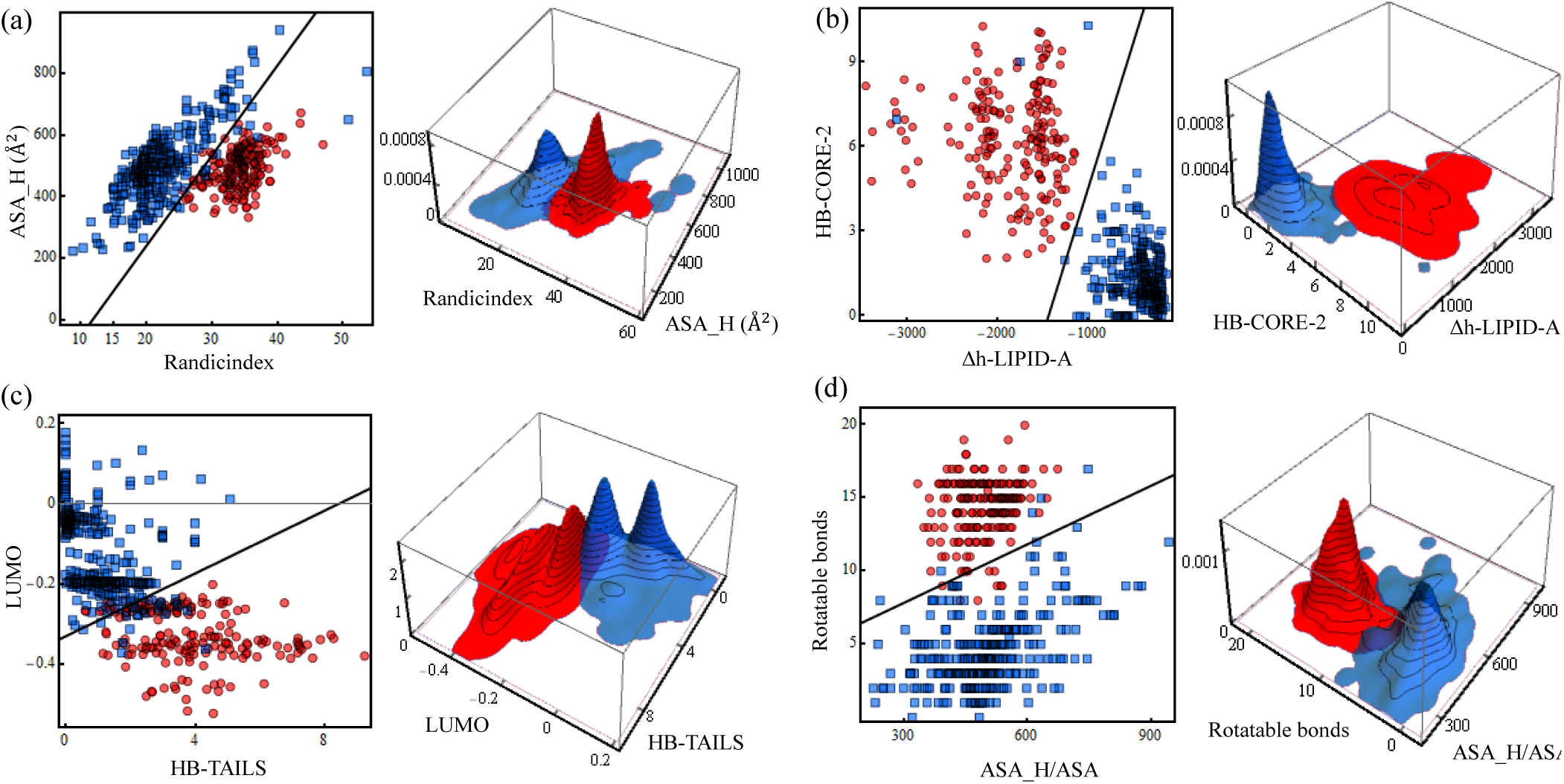
2-dimensional analysis of the parameter regions associated to weak (blue) and strong (red) permeation for four sets of highly accurate predictors found. Scatter plot at the left of each panel is equivalent to the density distribution plots at the right. Active compounds of the set G are shown on each panel. (a) Randicindex (QSAR), ASA H (Å^2^) (QSAR). (b) Δh-LIPID-A (permeation), HB-CORE-2 (permeation). (c) LUMO (QM), HB-TAILS (permeation). (d) Rotatable bonds (QSAR), ASA H/ASA (QSAR). Straight line in the left panel of each figure results from the analysis done by the two-dimensional SVM algorithm. The specifics of the function is given in Table S7.

**Figure S16:**
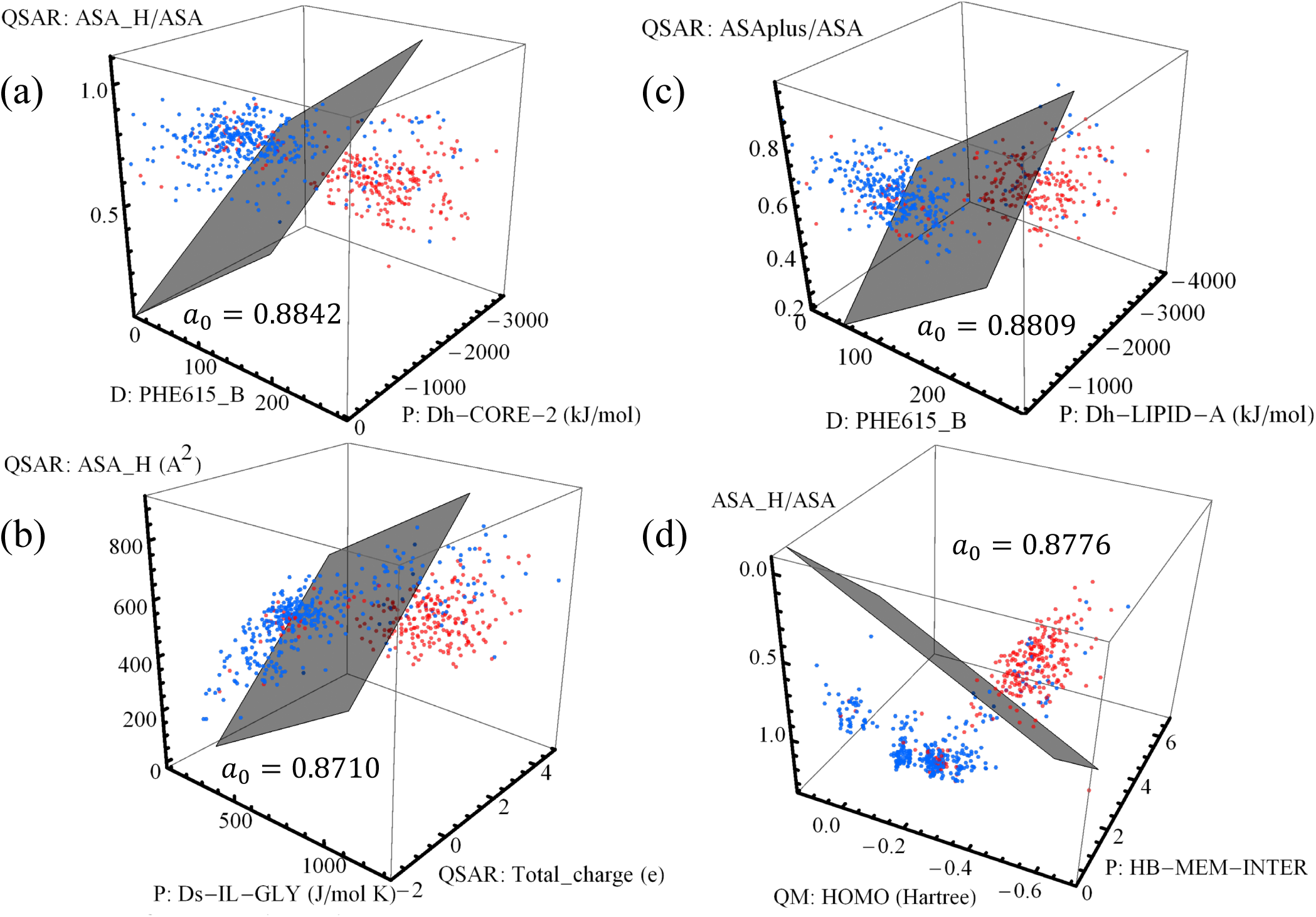
3-dimensional analysis of the parameter regions associated to weak (blue) and strong (red) permeation for four sets of highly accurate predictors found. All 605 active compounds are shown on each panel. (a) PHE615 B (docking), Δh-CORE-2 (permeation), ASA H/ASA (QSAR). (b) Δs-IL-GLY (permeation), Total charge (QSAR), ASA H(Å^2^)(QSAR). (c) PHE615 B (docking), Δh-LIPID-A (permeation), ASAplus/ASA (QSAR). (d) HOMO (QM), HB-MEM-INTER (permeation), ASA H/ASA (QSAR). Accuracy score *a*_0_ for each panel is also shown. Gray plane results from the analysis done by the three-dimensional SVM algorithm. The specifics of the function is given in Table S8.

**Table S7:**
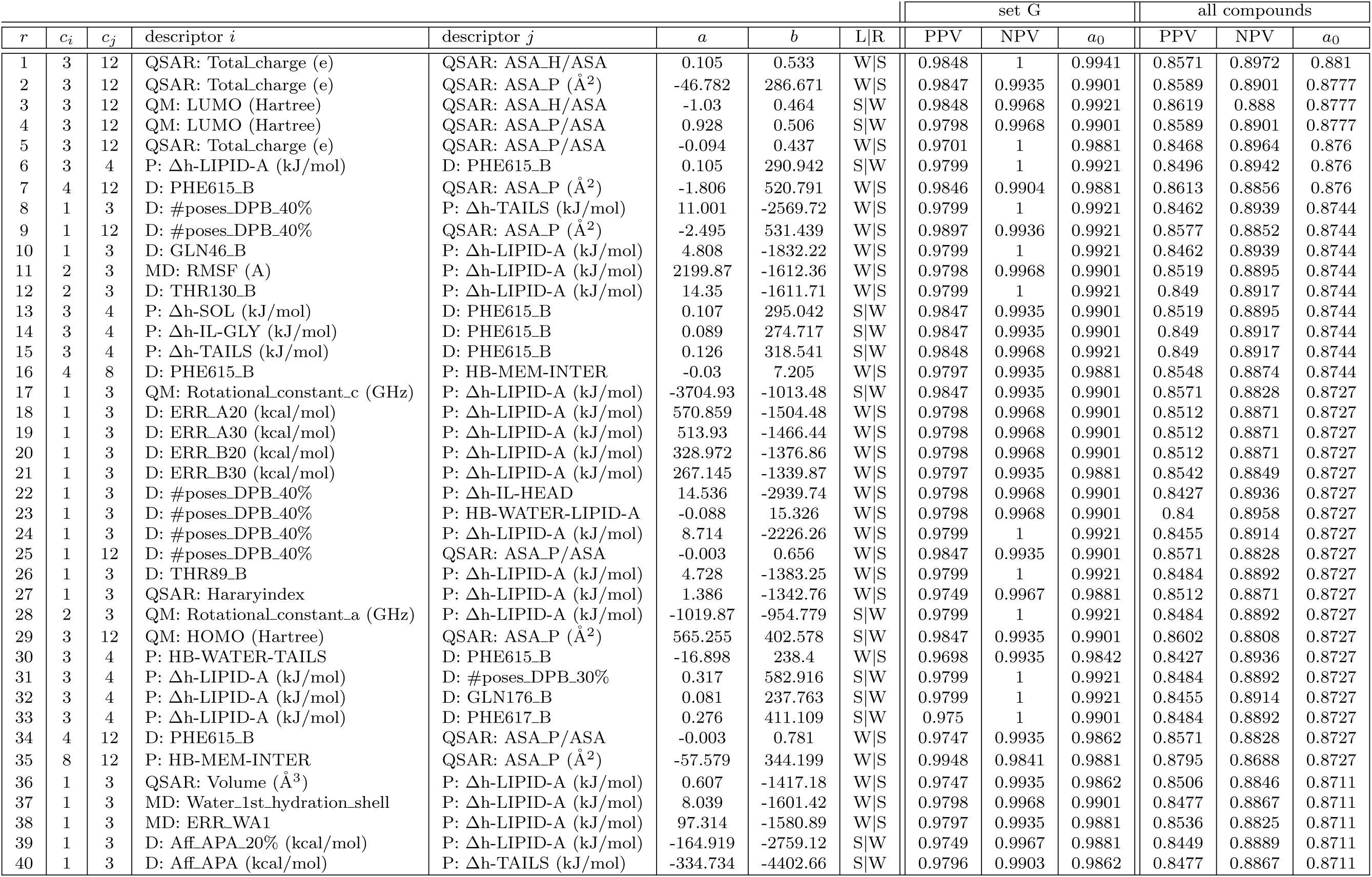
Top-40 pairs of descriptors and associated ranges with strong (SP) and weak (WP) OM permeation according to the accuracy score of all active compounds. The trained classification model over the compounds in the set G follow the linear equation *y* = *ax* + *b*, where *y* and *x* are the axes of descriptor *i* and *j*, respectively, and they are listed in the measurement units of the descriptors. Column L|R indicates whether WP are associated with descriptor values to the left of the trained linear model and hence SP associated with values to the right of the linear model (entry W|S), or vice versa (entry S|W). The evaluation metrics of positive predictive value (PPV), negative predictive value (NPV) and accuracy (*a*_0_) are listed for each descriptor for the compounds of the set G only and for all active compounds, as indicated.

**Table S8:**
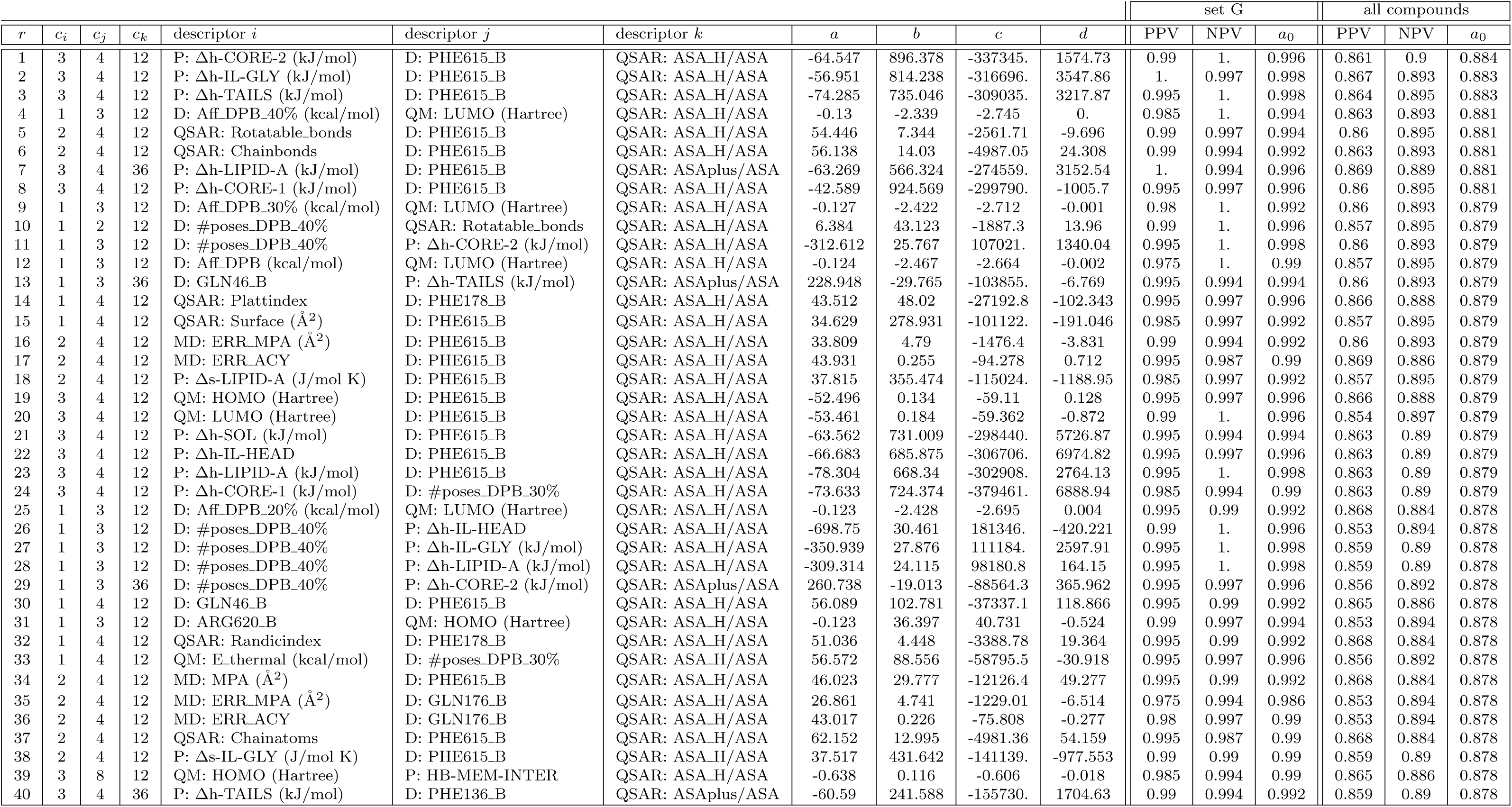
Top-40 groups of three descriptors and associated ranges with strong (SP) and weak (WP) OM permeation according to the accuracy score of all active compounds. The trained classification model over the compounds in the set G follow the equation *ax* + *by* + *cz* + *d* = 0, where *x*, *y*, and *z* are the axes of descriptors *i*, *j*, and *k*, respectively, and they are listed in the measurement units of the descriptors. The evaluation metrics of positive predictive value (PPV), negative predictive value (NPV) and accuracy (*a*_0_) are listed for each descriptor for the compounds of the set G only and for all active compounds, as indicated.

